# Identification of COVID-19 prognostic markers and therapeutic targets through meta-analysis and validation of Omics data from nasopharyngeal samples

**DOI:** 10.1101/2021.02.18.431825

**Authors:** Abhijith Biji, Oyahida Khatun, Shachee Swaraj, Rohan Narayan, Raju Rajmani, Rahila Sardar, Deepshikha Satish, Simran Mehta, Hima Bindhu, Madhumol Jeevan, Deepak K Saini, Amit Singh, Dinesh Gupta, Shashank Tripathi

## Abstract

While our battle with the COVID-19 pandemic continues, a multitude of Omics data has been generated from patient samples in various studies, which remains to be translated. We conducted a meta-analysis of published transcriptome and proteome profiles of nasal swab and bronchioalveolar lavage fluid (BALF) samples of COVID-19 patients, to shortlist high confidence upregulated host factors. Subsequently, mRNA overexpression of selected genes was validated in nasal swab/BALF samples from a cohort of COVID-19 positive/negative, symptomatic/asymptomatic individuals. Analysis of these data revealed S100 family genes (S100A6, S100A8, S100A9, and S100P) as prognostic markers of COVID-19 disease. Furthermore, Thioredoxin gene (TXN) was identified as a significant upregulated host factor in our overlap analysis. An FDA-approved drug Auranofin, which inhibits Thioredoxin reduction, was found to mitigate SARS-CoV-2 replication *in vitro* and *in vivo* in the hamster challenge model. Overall, this study translates COVID-19 host response Big Data into potential clinical interventions.

## INTRODUCTION

The COVID-19 pandemic has emerged as the biggest global public health crisis of this century. As of April 11, 2021, more than 136 million infections and 2.9 million casualties have been reported (https://www.worldometers.info/coronavirus/). The causative agent SARS-CoV-2 contains a single-stranded positive-sense RNA genome that encodes ∼27 proteins (1). COVID-19 disease is quite heterogeneous, and its manifestation ranges from asymptomatic, mild, severe to lethal, depending on a variety of host, virus, and environmental factors (2). Age, sex, ethnicity, and co-morbidities, all have been implicated in determining disease outcome (2-4). An effective and timely interferon (IFN) response is critical in resolving viral infections (3), however, SARS-CoV-2 has multiple strategies to suppress host immune response (4). Disruption of immune homeostasis and induction of cytokine storm has been recognized as one of the underlying causes of severe COVID-19 (5), yet the molecular mechanisms underlying immune dysregulation are yet to be defined.

Several research groups have applied tour de force high throughput methodologies to profile the host responses upon viral infections (6-13). This has resulted in a wealth of virus-host interaction Big Data, which hold the key to novel therapeutic strategies and molecular markers of infection and disease progression. Examination of host response at the primary site of infection in the upper respiratory tract is crucial to understand viral pathogenesis. Various studies have utilized BALF and nasopharyngeal swabs to characterize the changes in transcripts and proteins during infection to understand COVID-19 pathogenesis (6-12), which have highlighted significantly upregulated genes and biological pathways altered during infection. While proinflammatory cytokines, chemokines, enzymes in neutrophil-mediated immunity, and several IFN stimulated genes (ISGs) have consistently shown up in their analysis, experimental validation and mechanistic studies are generally lacking (7-12). A detailed characterization of antiviral responses in the upper respiratory tract of patients, its variation with age, sex, and association with progression of disease severity remains to be accomplished. (14-16).

The goal of our study was to identify genes that are consistently upregulated during SARS-CoV-2 infection in the upper respiratory tract of patients and understand their role in viral infection and disease progression. For this, we surveyed the literature for Omics data from COVID-19 positive patient’s nasal swab and BALF samples and selected 4 transcriptomic and 3 proteomic datasets. We performed a hypergeometric distribution-based overlap analysis followed by cumulative fold change score-based prioritization to shortlist genes. This was followed by an examination of selected gene expression levels in nasal swab/ BALF samples from a cohort of COVID positive, negative, symptomatic, and asymptomatic individuals, ranging from 30-60 years in age and of mixed gender. ROC analysis of gene expression data in nasal swabs revealed S100 family genes (S100A6, S100A8, S100A9, S100P) as high confidence markers of disease severity. Among other shortlisted genes, Thioredoxin (TXN) emerged as a significantly upregulated factor supported by multiple datasets. Thioredoxin is a proinflammatory protein that requires to be reduced by Thioredoxin reductase enzyme, which itself can be targeted by an FDA-approved gold drug Auranofin (17, 18). We tested the antiviral efficacy of Auranofin in cell culture and preclinical Syrian hamster challenge model and found that it can reduce SAR-CoV-2 replication over 1 order of magnitude at a well-tolerated non-toxic dosage. This drug is already in clinical use for inflammatory diseases and can be considered for COVID-19 treatment based on our data.

Through collective global efforts several COVID-19 vaccines have become available in an astonishingly short period, although new virus variants have emerged, some of which can escape vaccine-mediated immunity (19). Progress on the development of the antivirals and disease prognostic markers has been lagging. Repurposing clinically approved drugs for use against SARS-CoV-2 has been an attractive option and has been explored by many research groups through different approaches (20). Our study translates COVID-19 virus-host interaction and response Big Data into potential actionable clinical interventions, including the use of S100 genes as a prognostic marker in the nasal swabs and repurposing clinically approved drug Auranofin for COVID-19 treatment.

## RESULTS

### Compilation and overlap analysis of published transcriptomics and proteomics data from COVID-19 patient samples revealed 567 upregulated host factors

We started the study by compiling the host factors that are consistently and significantly upregulated in the upper respiratory tract of COVID-19 patients. For this, we decided to use published transcriptomics and proteomics datasets derived from nasal swab or BALF samples of COVID-19 patients. We chose four transcriptomics (T), and three proteomics (P) datasets and further analysis was performed according to a rationally designed workflow (Figure 1A). All datasets included differentially expressed genes in infected patients with healthy individuals as control (Table S1). The selection criteria (described in materials and methods) included at least 1.5-fold (2-fold for one dataset) gene upregulation at both mRNA and protein levels. The filtration of data was carried out to sort only significantly upregulated genes from all the datasets (Table S2). A pairwise overlap analysis was performed on the filtered genes/proteins from each study and significantly overlapping genes (p-value < 0.01 calculated using Fisher’s exact test) between T1-T3 (14), T1-T4 (9), T1-P3 (2), T3-T4 (504), T3-P1 (10), T3-P2 (8), T3-P3 (17), T4-P1 (8). T4-P3 (15) and P1-P3 (3) were determined (Figure 1B, Supplementary File 1). This method was adapted from similar overlap analysis conducted previously to compare multiple virus-host interaction datasets and obtain the significance of intersections (21). Union of intersections between the T-T and T-P and P-P after the overlap analysis results in 567 genes (Figure 1B). To reiterate the functional characteristics of the differentially expressed genes (DEGs), we examined the biological processes and signaling pathways they are involved in. Pathway enrichment of 567 genes from the union of all intersections from overlap analysis (TT+TP+PP) showed enrichment of biological processes like protein elongation, interferon (IFN) signaling, chemotaxis of granulocytes, and inflammatory pathways (Figure 1C). The antiviral response to respiratory viral infections including SARS-CoV-2 is driven by interferons (IFNs) (16). Hence, we examined the shortlisted set of genes for their potential regulation by different categories of IFNs, using the Interferome tool (22). We found that out of 567 genes, 205 were regulated by type I IFN, 170 genes by Type II IFN, 327 genes were regulated by both type I and type II IFN, while 16 genes were commonly regulated by all the three classes of IFNs (Figure 1D). These 16 genes are well-characterized interferon-stimulated genes (ISGs), that include direct antiviral effector ISGs (IFITs, MX1, OAS3, and OAS1), as well as positive regulators (STAT1) of IFN response (23). This indicated an active IFN mediated innate antiviral response in the upper respiratory tract cells during SARS-CoV-2 infection and highlighted potential antiviral factors.

**Figure 1:**
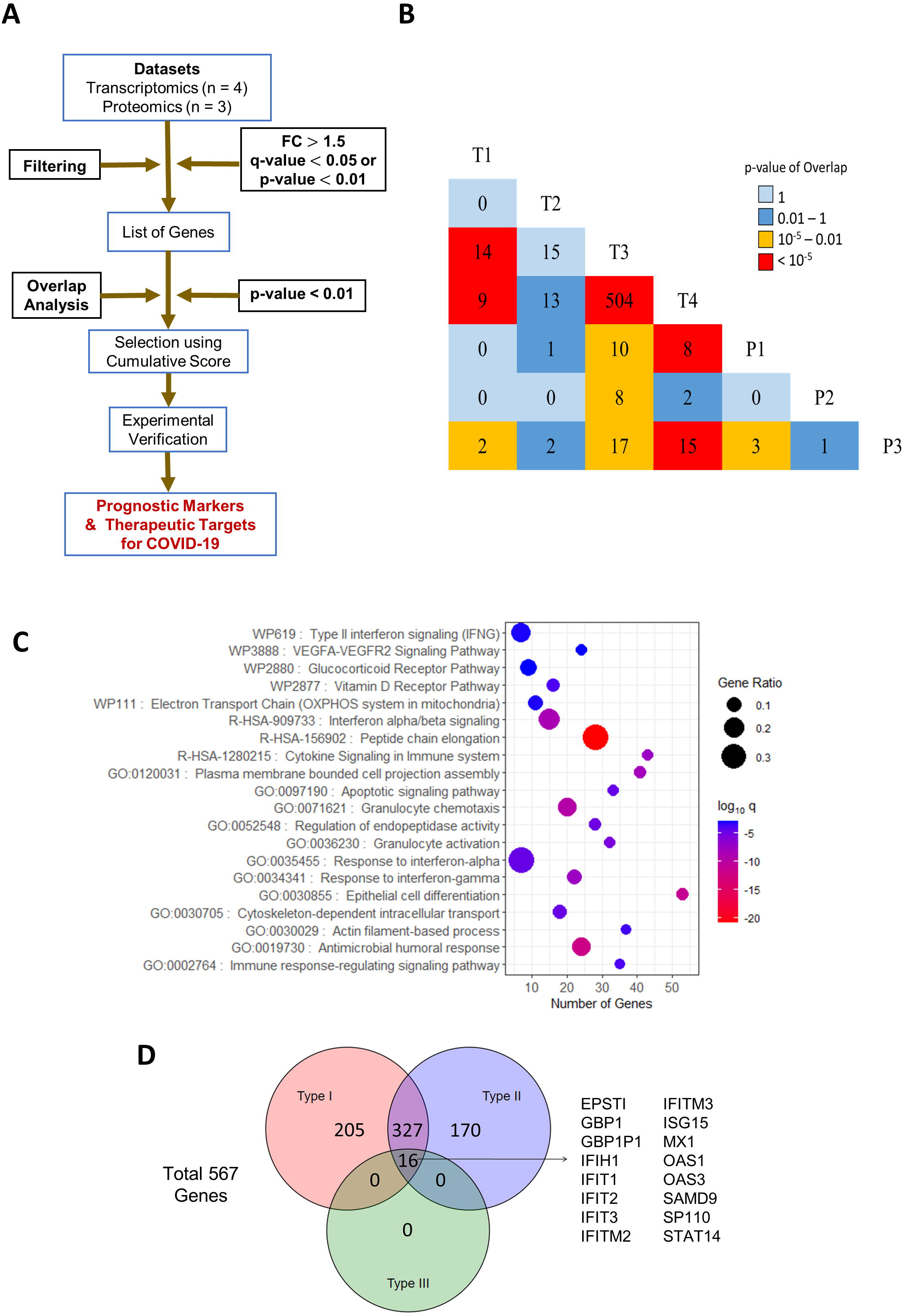
Meta-analysis pipeline for gene prioritization and associated pathway analysis. **A)** Three proteomics and four transcriptomics datasets were chosen to obtain biomarkers for COVID-19 in humans. Genes/proteins that came up in these studies with a fold change greater than 1.5 and a q-value less than 0.05 (p-value less than 0.01 was taken in cases where q value is not provided) were subjected to pairwise overlap analysis. Genes that fall under significant intersections and represented in at least one proteomic dataset were sorted using cumulative scores to be experimentally verified. **B)** Triangular heatmap showing pairwise overlaps between transcriptomic and proteomic datasets. The number within each box denotes the number of genes that showed up between the corresponding intersections. The color of a box denotes the significance of overlap determined by Fisher’s exact test. **C)** Gene ontology of all genes (567) in the significant intersections obtained during the overlap analysis plotted with the number of genes in each term on the X-axis, proportion of genes enriched compared to the total number of genes in each term as the size of dots and the color representing log_10_ p-adj value (q-value) of enrichment. **D)** Venn diagram showing the number of genes that are induced by Type I, II, or III interferons. The analysis was performed on Interferome v2.01 using the union of significant intersections (567)

### Rank ordering and shortlisting of upregulated host factors highlighted host factors regulating the antiviral and inflammatory immune response in COVID-19 patients

Since proteome dictates the outcome inside a cell, soluble factors are key in shaping the antiviral response. We focused on genes supported by orthogonal transcript (T) and protein (P) abundance data. For this, we chose genes from the union of intersections of T-T, T-P, and P-P overlaps, which was reported at least in one of the proteomics studies. This narrowed down the list to a total of 46 genes that were intersecting in T-P (26), P-P (2), TT-TP (16), TP-PP (1), and TT-TP-PP (1) overlaps (Figure 2A and 2B). A cumulative score for the 46 selected significantly upregulated genes was calculated using the sum of their log_2_ fold-change values in the parent datasets and ranked (Figure 2C). The enrichment of these 46 genes in each of the datasets, where the expression is reported, is shown in Figure 2B. Many of these genes are directly regulated by different classes of interferons. 15 genes are regulated by IFN-I, while 8 genes by IFN-II. 20 genes are regulated by both type-I and type-II IFNs, while only 2 genes by all the three types of IFNs (Figure 2D). Most of the IFITs and other ISGs that were earlier determined in our analysis to be regulated by all the three types IFNs are no more in the list since those ISGs were only reported upregulated at transcriptome level (only in T-T overlap) and hence were lost when the genes were filtered for their upregulation at the protein level, leaving behind only MX1 and OAS3 (Figure 1C and 2D). The biological functions of the selected 46 genes were also investigated to understand their roles in COVID-19 pathophysiology. The pathways enriched were mainly related to innate immune response and defense against microbes along with inflammatory and immune signaling, neutrophil degranulation, and cellular response to TNF and interferon-gamma (Figure 2E).

**Figure 2:**
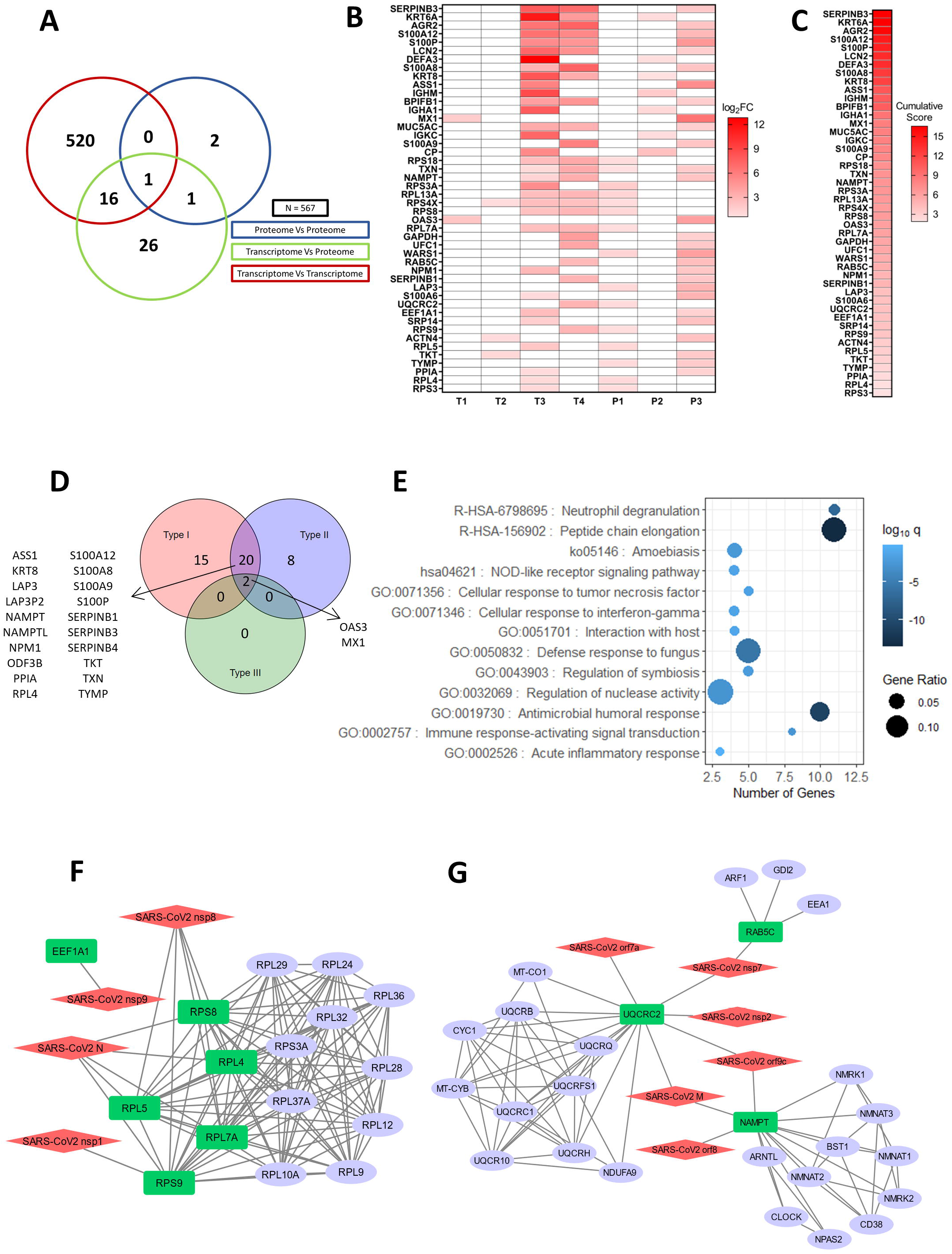
Cumulative score ranking, pathway, and interactome analysis of selected host factors. **A)** Venn diagram of genes obtained from significant intersections among proteomic or transcriptomic datasets after pairwise overlap analysis. **B)** Genes in the Venn diagram that were found in at least one proteomic dataset with their log_2_FC values in the datasets where they are present. Boxes colored in white denote that the gene is not present in the particular dataset. **C)** Genes arranged in descending order of cumulative scores obtained as a sum of log_2_FC values in the datasets where they are present. **D)** Venn diagram showing the number of interferon-induced genes performed using Interferome v2.01 for 46 selected genes. **E)** Gene ontology of 46 genes plotted with the number of genes in each term on the X-axis, the proportion of genes enriched compared to the total number of genes in each term as the size of dots and the color representing log_10_ p-adj value (q-value) of enrichment. **F, G)** Virus-host protein-protein interactions among SARS-CoV2 proteins and significant genes in the overlap analysis that shows up in at least one proteomic dataset modeled using Cytoscape v3.8.0. A STRING interactome for the primary interactors of SARS-CoV-2 proteins was merged (confidence ≥0.999 for all the proteins and confidence ≥0.90 for NAMPT; max number of interactors = 10). Red: SARS-CoV-2 proteins, Green: Host proteins (primary interactor), blue: STRING interactors (other cellular proteins interacting with the primary interactors).

Further, to understand the potential role of shortlisted genes in COVID-19 pathophysiology, their interactions with SARS-CoV-2 proteins were inspected by analyzing the publicly available SARS-CoV-2 cellular interactome data (24). For this, host protein-protein interactions were retrieved from the STRING database (25) and merged with the virus-host protein-protein interactions giving a discrete picture of how the viral proteins target various cellular processes during infection. Other than NAMPT, UQCRC2, and RAB5C, it was mainly ribosomal proteins that were primary interactors to the SARS-CoV-2 proteins (Figure 2F and 2G). We also examined the intracellular, cellular, tissue, and organ-specific expression for shortlisted genes using publicly available data (26). Many upregulated proteins were predicted to localize in the intracellular organelles like endoplasmic reticulum, mitochondria, Golgi complex, and endosomes (Figure S1A), while 19 genes were predicted to be secretory. A thorough analysis of the list of 46 selected genes using Human Tissue Atlas revealed that they are expressed in the respiratory tract and in immune effector cells known to survey infection sites (Figure S1B). The relative expression levels show that genes associated with protein synthesis (ribosomal proteins and elongation factors) are highly expressed compared to any other genes and are enriched across all the tissues in the map (Figure S1B).

### qRT-PCR based validation in a cohort of COVID-19 positive/negative, symptomatic/asymptomatic individuals reveals differential upregulation of selected genes in a disease-specific manner

For validation using qRT-PCR and further analysis, we selected genes with a cumulative score greater than 10 (Figure 2C). Also, we considered genes belonging to the S100 family that came up within 46 shortlisted genes, since they are known regulators of inflammation (27, 28). Furthermore, we also selected the TXN since it was supported by multiple lines of evidence and appeared in the TT-TP-PP overlap in our study (Figure 2A). The COVID-19 patient cohort used for qRT-PCR of genes, included 63 individuals (both males and females, aged 30-60 years), out of which 16 each were COVID-19 positive-symptomatic (PS), COVID-19 positive-asymptomatic (PA), COVID-19 negative-symptomatic (NS), and 15 were COVID-19 negative-asymptomatic (NA) healthy category (Table 1). Total mRNA from the nasal swab was isolated and the upregulation of 14 selected genes was verified by qRT-PCR. The log_2_ fold-change expression with respect to the average of the negative asymptomatic group (Figure S2, Figure 3A) was calculated and plotted on the heatmap (Figure 3A), which depicts the mRNA enrichment of the selected genes in different patient samples and categories. Next, we determined the correlation between the viral RNA load in COVID-19 patients (qRT-PCR of viral envelope (E) gene) and log_2_ Fold-change of selected host genes in the patient’s sample. It was observed that the Ct value for the E gene was negatively correlated with log_2_ Fold-change of genes showing that viral load and disease severity are positively correlated (Figure S3). Furthermore, the upregulation of selected host genes was more pronounced in positive symptomatic patients with a higher viral load than positive asymptomatic individuals (Figure 3A and Figure S3). A comparative heatmap in Figure 3B gives an insight into the genes that can be considered as COVID-19 disease and/or severity marker. While all the upregulated genes indicate infection (Figure 3B; NA-PS), only a few genes showed significant upregulation in a COVID-19 specific manner (Figure 3B; NS-PS).

**Table 1:** Summary of individual and different categories in the COVID-19 cohort used for qRT-PCR based validation analysis. All samples were collected from Bangalore Urban area for diagnostic purposes.

| Patient Status | Number of patients | Average age | Number of males | Number of females | Number in the age group 30-40 | Number in the age group 41-50 | Number in the age group 51-60 |
| --- | --- | --- | --- | --- | --- | --- | --- |
| Negative Asymptomatic | 16 | 43.9 | 8 | 8 | 5 | 6 | 5 |
| Negative Symptomatic | 16 | 41.7 | 12 | 4 | 9 | 4 | 3 |
| Positive Asymptomatic | 15 | 44.3 | 8 | 8 | 6 | 6 | 4 |
| Positive Symptomatic | 16 | 45 | 8 | 8 | 5 | 5 | 6 |

**Figure 3:**
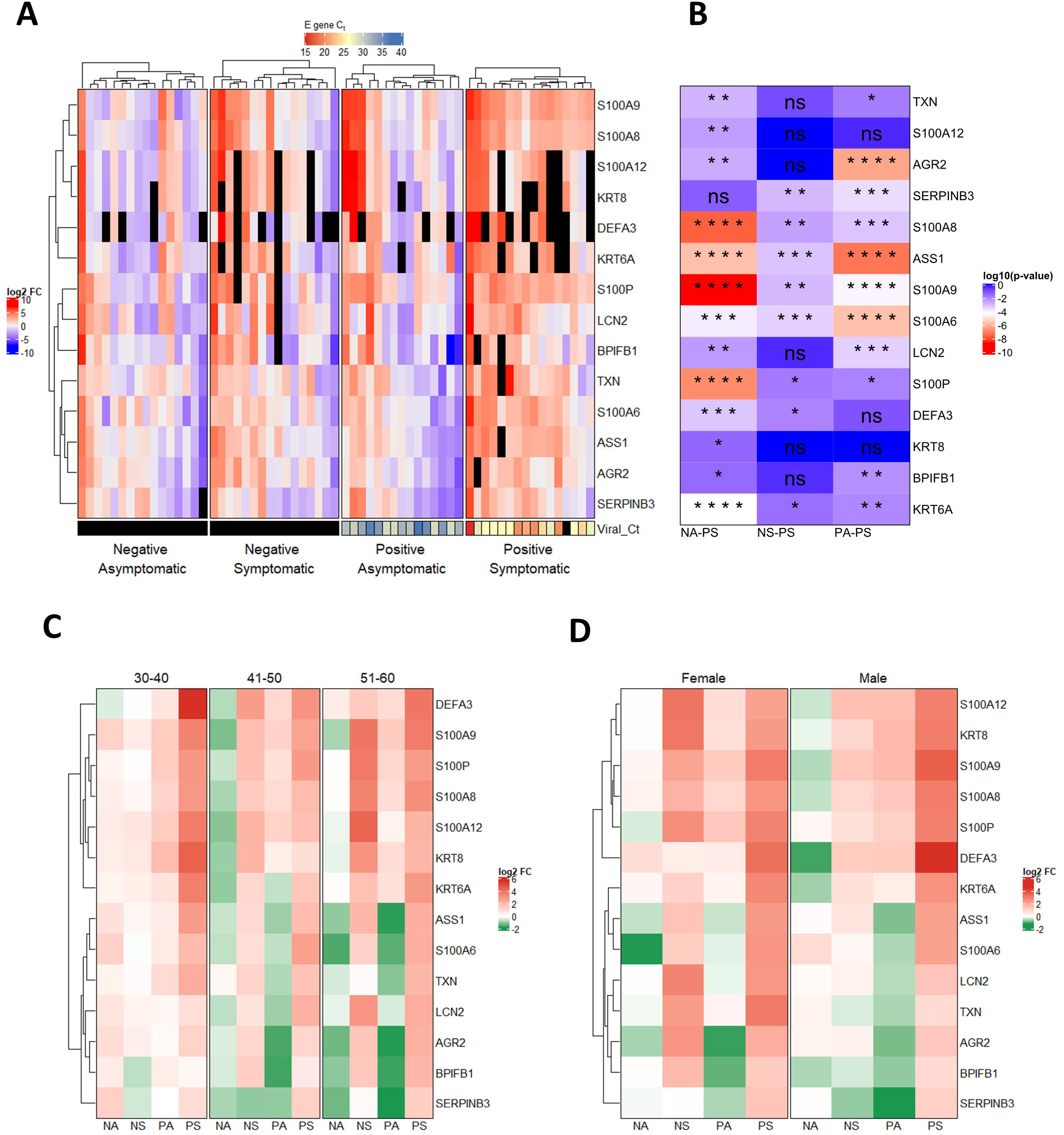
qRT-PCR validated expression profile of selected genes in different categories of COVID-19 cohort. A) qRT-PCR was performed on RNA isolated from COVID-19 patients for 14 genes and average log_2_ Fold-change values (with respect to Negative Asymptomatic group) of PCR triplicates are shown in a heatmap. Each column represents a patient and clustering was performed for columns and within row slices. The bottom annotation shows the C_t_ value for the viral gene encoding Envelope (E) protein with a corresponding legend on the top. Black boxes denote ‘value unknown/undetermined. **B)** Differences between groups for each gene were computed using the Kruskal-Wallis test followed by post hoc Dunn’s test with Bonferroni corrections for multiple comparisons. The log_10_ (p-value) of comparisons is shown in the heatmap. The comparisons are Negative asymptomatic vs Positive symptomatic (NA-PS), Negative symptomatic vs Positive symptomatic (NS-PS), and Positive asymptomatic vs Positive symptomatic (PA-PS). *P < 0.05; **P < 0.01; ***P < 0.001; ****P < 0.0001; ns – not significant. **C)** log_2_ Fold-change values are grouped based on age groups 30-40, 41-50, and 51-60. Each row represents the average of log_2_ Fold-change values for patients falling into the particular age group and respective disease status. **D)** log_2_ Fold-change values are grouped according to sex. Each row represents the average of log_2_ Fold-change values for patients falling into the particular sex and respective disease status.

Multiple genes from the S100 family, including S100A8, S100A9, S100A6, and S100P, and few other genes such as ASS1 and SERPINB3 were significantly upregulated in positive symptomatic patients when compared to other three categories (NA, NS, PA), suggesting their potential diagnostic and prognostic value (Figure 3B, NS-PS). Expression of neutrophil defensin alpha 3 (DEFA3) was upregulated in some of the positive symptomatic patients but remained undetermined in many cases. Furthermore, we examined the influence of age and sex on the upregulation of selected gene in patient’s samples by categorizing them based on age groups (30-40, 41-50, and 51-60) and gender (male and female) (Figure 3C, Figure 3D, Figure S4 and S5). The qRT-PCR data revealed that all the selected genes were induced in positive symptomatic patients, irrespective of age or gender. However, closer examination of the heatmap reveals S100 family genes (S100A8, S100A9, S100P) being upregulated to a higher level in the 30-40 year age group and male individuals (Figure 3C, 3D).

### ROC analysis of mRNA expression of shortlisted significant genes in the COVID-19 cohort unveils the prognostic potential of the S100 family of genes

The COVID-19 symptomatic group of patients included individuals with breathing difficulty, fever, hospitalization, and SARI (severe acute respiratory infections), whereas asymptomatic patients had none of these features (Table1). To evaluate the prognostic value of selected genes in differentiating asymptomatic vs symptomatic COVID-19 cases, we conducted a non-parametric Receiver Operating Characteristic (ROC) curve analysis (29) for the 11 genes that were significant after comparison between positive symptomatic and asymptomatic group (Figure 3B). For this, we used their threshold cycle (C_t_) values for COVID-19 positive cases to plot the curve, and the area under the curve (AUC) was computed (Figure 4A). All genes were found to significantly differ (AUC > 0.5) from the line where True positive rate = False positive rate, indicating their potential to differentiate between asymptomatic and symptomatic individuals (Figure 4B). The optimal C_t_ value cut-off was determined for significant genes using the ROC01 method which finds the point in the ROC curve closest to (0,1) corresponding to 100% specificity and sensitivity. Since the prognostic marker should correctly identify symptomatic patients from asymptomatic ones, we looked at the genes with maximum sensitivity while not compromising on specificity at the optimal cut-off. S100A8 (Cut-off = 9.964663, Sensitivity = 0.938, Specificity = 0.688) had the highest sensitivity at the optimal cut-off. Other S100 family members like S100A9 (Cut-off = 8.533607, Sensitivity = 0.854, Specificity = 0.729), S100A6 (Cut-off =8.472503, Sensitivity = 0.745, Specificity = 0.718) and S100P (Cut-off = 11.23458, Sensitivity = 0.812, Specificity = 0.622) also showed good prognostic potential (Figure 4C and 4D). Genes like LCN2 (Cut-off = 11.23362, Sensitivity = 0.744, Specificity = 0.756), AGR2 (Cut-off = 11.19266, Sensitivity = 0.775, Specificity = 0.708) and ASS1 (Cut-off = 12.70913, Sensitivity = 0.7, Specificity =0.771) were also found to have desired sensitivity and specificity values (Figure S6).

**Figure 4:**
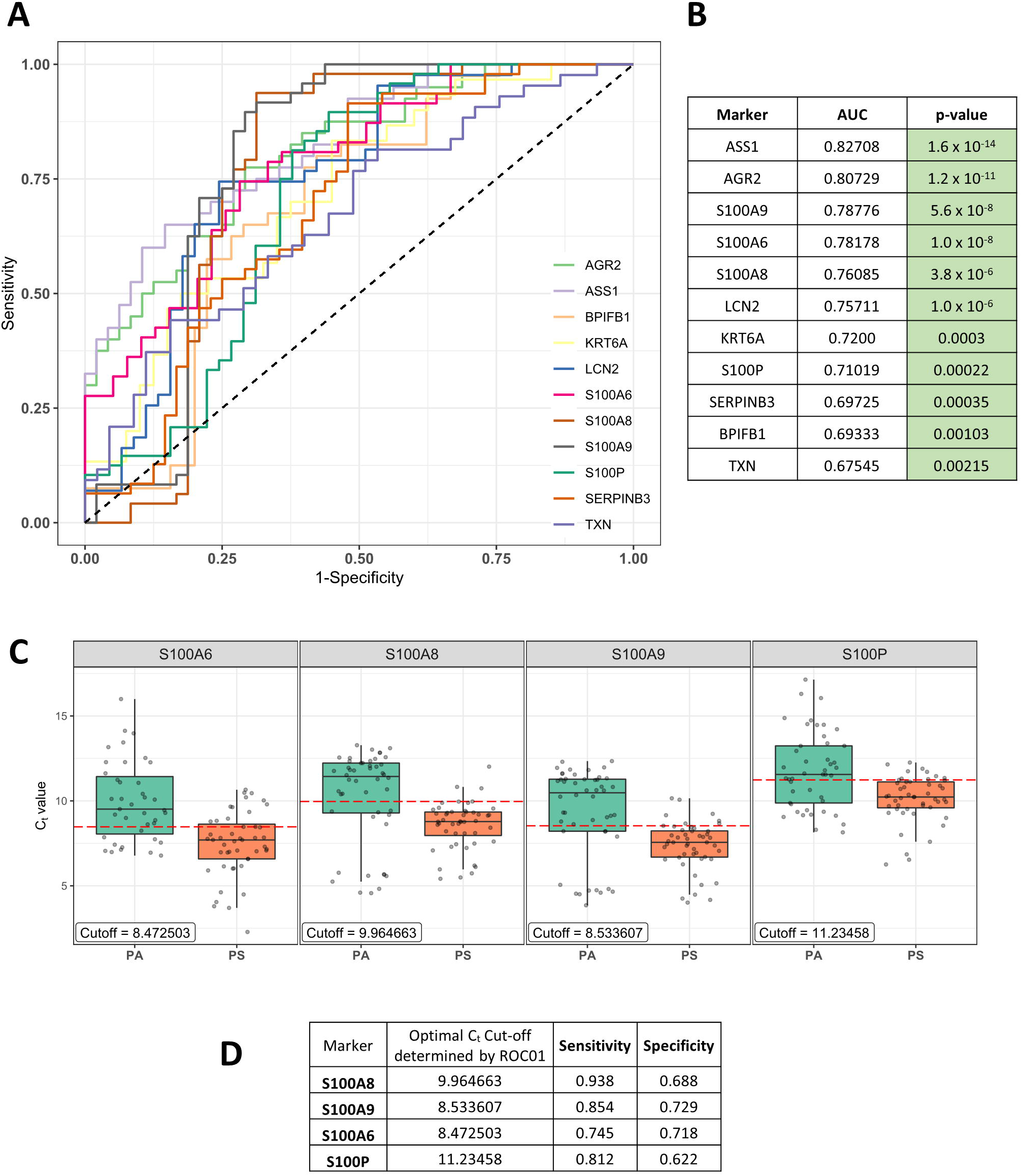
ROC analysis of genes in COVID-19 positive patients to identify prognostic markers. **A)** ROC curve for C_t_ value of genes in COVID-19 positive patients. The black dashed line corresponds to no prognostic potential where True positive rate (Sensitivity) and False positive rate (1-Specificity) are equal. **B)** The value of Area Under the Curve (AUC) for each ROC curve along with the p-value. Significance was calculated non-parametrically (DeLong’s estimate) using the Wald test statistic. **C)** Boxplot of C_t_ values for significant S100 family of genes in Positive asymptomatic (PA) and Positive symptomatic (PS) patients. The red dashed line shows the optimal C_t_ cut-off determined by the ROC01 method (also shown in the label in each graph). **D)** Optimal C_t_ cut-off, sensitivity, and specificity values for significant S100 family of genes.

### Thioredoxin reductase inhibitor drug Auranofin significantly mitigates SARS-CoV-2 replication *in vitro* and *in vivo* in the hamster challenge model

Thioredoxin (TXN) was a single hit that appeared in the TT-TP-PP overlap in our study and remained in the shortlisted gene set at the end of the meta-analysis. Although its expression upregulation or the prognostic value was not the highest, it is part of a druggable pathway. Thioredoxin is known to promote inflammatory cytokine induction, apoptosis, and regulate redox status, for which it switches between oxidized and reduced forms through the action of thioredoxin reductase, which can be inhibited by an FDA approved orphan drug Auranofin (2,3,4,6-tetra-o-acetyl-L-thio-β-D-glycopyranp-sato-S-(triethyl-phosphine)-gold) (30, 31). We sought to check the effect of Auranofin, which will lock Thioredoxin in its oxidized form, on SARS-CoV-2 infection and replication in cell culture and animal models. To begin, cell viability assay performed in HEK-ACE2 cells using increasing doses of Auranofin showed minimal cytotoxicity at the lowest concentration (1µM) and had predicted CC_50_ of 9.659µM (Figure 5 A). The effects of increasing doses of Auranofin up to 1 µM, was then tested on SARS-CoV2 replication *in vitro*. For this, cells were pretreated with the drug which remained present during the course of infection. Analysis of viral RNA 48hr post infection showed a reduction of more than one order of magnitude, starting at treatment with 0.25 µM Auranofin (Figure 5B). With a calculated EC_50_ = 0.29µM, the selectivity index (CC_50_/IC_50_) of auronafonin was determined to be 33.3. The potent antiviral effect of Auranofin was confirmed by western blot for the full-length viral spike protein (Figure 5 C). Next, we decided to confirm the antiviral activity of Auranofin in Syrian golden hamsters, which are currently considered as the animal model of choice to evaluate vaccines and antivirals (32). Auranofin (PubChem CID 6333901) toxicity and bioavailability in rodents have been described before (33), based on which we first tested its oral toxicity in hamsters at 1mg and 5mg/kg body weight, which showed the drug was well tolerated at the tested doses (Figure S7). For infection studies, the drug was orally administered in prophylactic and therapeutic formats; before and after infection respectively (Figure 5D). The viral titers in lungs of animals at Day 4 revealed that therapeutic administration of Auranofin with a non-toxic concentration of 5mg/kg body weight was more effective at mitigating virus replication (reduction by more than one order of magnitude) in lung tissue, compared to prophylactic dosage (Figure 5E). Bodyweight loss results were also indicative of the same when compared to the virus challenge group (Figure 5F). Also, we found that the TXN gene was upregulated in the lungs of infected animals compared to the mock group, which correlates to our findings from patient sample gene expression data (Figure S8)

**Figure 5:**
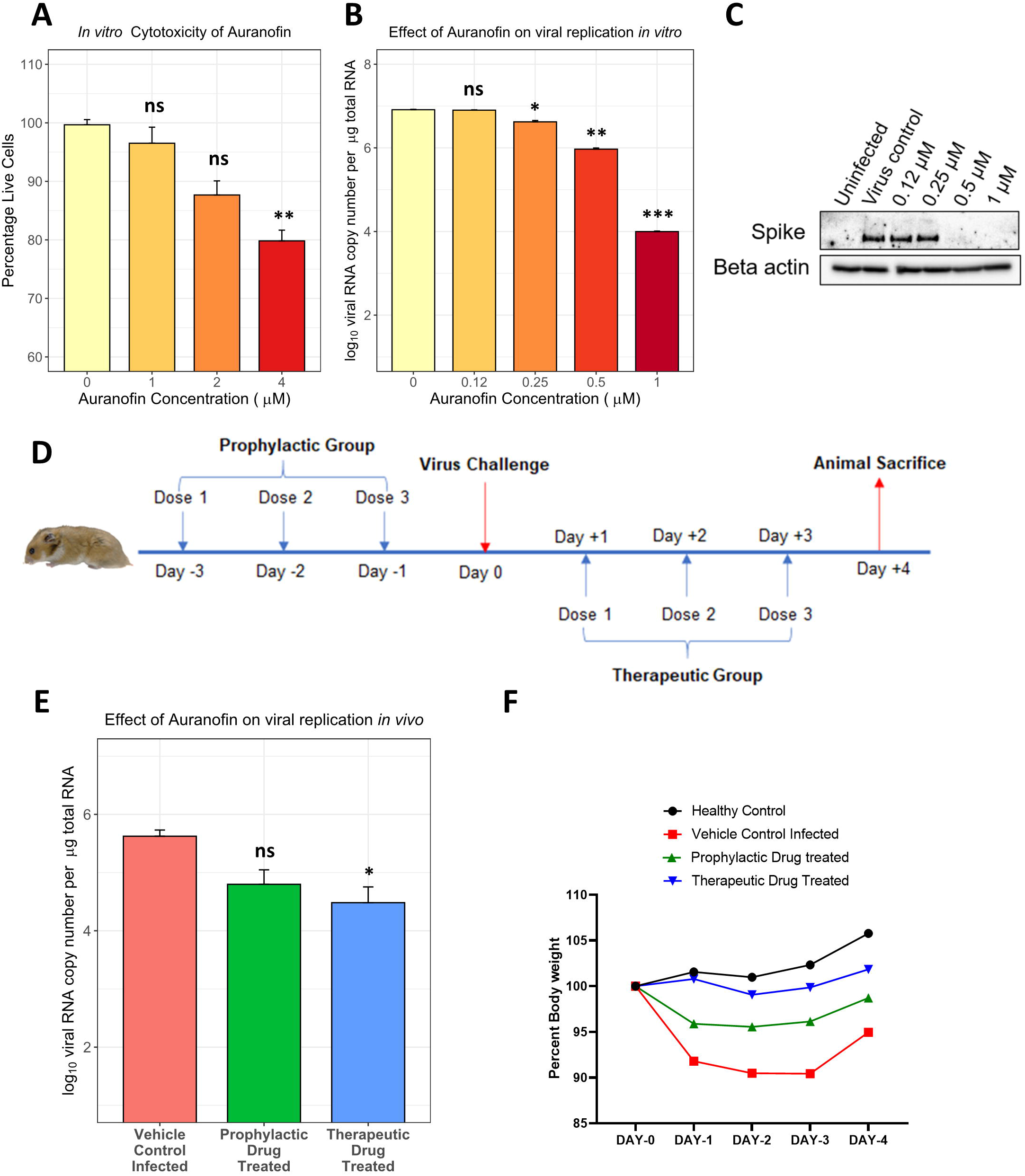
Auranofin inhibits SARS-CoV-2 replication in cell culture and preclinical hamster challenge model. **A)** HEK-ACE2 cells were treated with the indicated dose of the drug for 48 hours. Cell viability was measured and plotted on the graph. **B)** HEK-ACE2 cells were pre-treated with the indicated amount of drug for 3 hours and then infected with SARS-CoV-2 at 0.1 MOI for 48 hours. Total RNA was isolated from the cells and viral RNA copy number was measured by qRT-PCR. Log_10_ copy number of vRNA per µg total RNA is plotted on the graph. **C)** HEK-ACE2 cells were pre-treated with the indicated amount of drug for 3 hours and then infected with SARS-CoV-2 at 0.1 MOI for 48 hours. Cells were harvested with 1x Laemmli buffer and probed for spike and beta-actin. **D)** 10–12-week-old hamsters (4 per group) were pre-treated with 5mg/kg bodyweight drug through oral route in 200 µl PBS vehicle. For the prophylactic group, this was done once per day for 3 days before infection, and for the therapeutic group, it was done once per day for 3 days post-infection. SARS-CoV-2 inoculum (10^5^ pfu/100 µl) was administered intranasally. The vehicle control group was administered the corresponding volume of DMSO in PBS same as the prophylactic group. Day 4 post-infection animals were sacrificed to measure viral and cellular RNA quantity in lung tissue. **E)** Total RNA was isolated from the lung tissue of infected animals and viral RNA copy number was measured by qRT-PCR. Log10 copy number of vRNA per µg total RNA is plotted on the graph (n=4). **F)** Body weight of hamsters was measured from D0 to D4 and plotted on the graph, considering weight on D0 as 100% (n=4). Error bars represent mean + standard error. Differences between test groups and control group were computed using the t-test with Bonferroni corrections for multiple comparisons. *P < 0.05; **P < 0.01; ***P < 0.001; ****P < 0.0001; ns – not significant.

## DISCUSSION

Several studies have analyzed changes in global transcriptome and proteome in COVID-19 patient samples of various kinds (6-12). These studies have given an overview of the biological processes that are modulated during SARS-CoV-2 infection; however, translation of this knowledge into antiviral interventions requires validation and mechanistic studies. Meta-analysis of virus-host interaction Big Data is a useful approach to narrow down key host factors and processes involved in viral replication and pathogenesis (21, 34). In our study, we focussed on transcriptomics and proteomics data from COVID-19 positive nasal swab and BALF samples and performed an integrative analysis to identify host factors involved in SARS-CoV-2 infection and disease progression. We reasoned that changes at mRNA levels must also be manifested at the protein level to bring out phenotypic differences in the infected individuals. Hence, we designed our meta-analysis pipeline to shortlist genes that were represented in orthogonal transcriptomics as well as proteomics datasets. Expression of the genes selected through meta-analysis was examined in nasal swab/BALF samples collected for COVID-19 diagnosis from a cohort of individuals that were COVID-19 negative or positive, and within those two categories either symptomatic or asymptomatic. The cohort design was to ensure the identification of genes that are overexpressed in a COVID-19 specific manner and those which indicate disease severity. The initial compilation of upregulated factors had 567 genes, of which 46 genes passed through the selection pipeline (Figure 2B). Most of these genes turned out to be IFN regulated and among them, the major category was ribosomal proteins (RPs) including RSP3A, RPL4, RPL5, RPL18, RPL13A, RPS4X, RPL7A, RPS9, and RPS3 (Figure 2B). RPs have been reported to be hijacked by different viruses, including SARS-CoV-2 during infection to shut off host translation and facilitate IRES-mediated translation of viral proteins (35-37). Inspection for reported interactions between shortlisted RPs with the SARS-CoV-2 proteins revealed that nsp1, nsp8, nsp9, and nucleocapsid (N) proteins of SARS-CoV-2 are potential interactors (Figure 2F). This suggests extensive targeting of host translational machinery by multiple SARS-CoV-2 proteins in the upper respiratory tract cells. Other shortlisted cellular proteins with reported interactions with viral proteins were NAMPT, UQCRC2, and RAB5C (Figure 2G). These are involved in ATP production, NAD synthesis, and vesicular fusion respectively, all of which have been reported to be modulated during SARS-CoV-2 infection (38, 39).

Subsequent ranking of genes based on cumulative upregulation score across different datasets, with dual support from transcriptomic and proteomic evidence, shortlisted 14 high confidence upregulated genes (Fig 2B). To confirm their upregulation during SARS-CoV-2 infection and the effect of patient age, sex, disease severity on the same, their expression was measured in a cohort of patients described earlier (Table 1). The data revealed that 11 genes were upregulated significantly in the PS category when compared to PA and hence had prognostic value. Whereas 8 genes were upregulated when compared to the NS category, hence had diagnostic value (Figure 3B). The data indicated higher levels of selected gene expression in younger male patients, which is consistent with previous reports of age and sex-dependent differences in COVID-19 induced gene expression and disease severity (6, 40). Among host factors that appeared at the end of meta-analysis and validation in the COVID-19 cohort, the S100 family of genes (S100A6, S100A8, S100A9, S100A12, S100P) emerged as a major group. An upregulation of S100 proteins is reported previously as an indication of viral or bacterial infections (27). The extracellularly secreted S100 proteins include S100A12, S100A8, and S100A9 (Figure S1A), all of which have been shown to serve as a danger signal and in regulating immune response (28). They activate NF-kB signaling through RAGE and TLR4 pathways stimulating the cells to produce proinflammatory cytokines at the site of infection (28). Several studies have explored serum diagnostic and prognostic markers by evaluating transcriptomic and proteomic changes in mild, severe, and fatal cases of COVID-19 (41, 42). An increase in S100A8/A9 (calprotectin) levels in serum have been correlated with severe forms of the disease (43). Transcriptomic studies on lung tissue of fatal COVID-19 cases have also reported which report an upregulation in S100A12, S100A8, S100A9, and S100P in patients (44, 45). In our study, the ROC01 curve analysis of the PA and PS group qRT-PCR data showed that all shortlisted S100s (except S100A12) had significant sensitivity as a prognostic marker of symptomatic COVID-19 (Figure 4 C, D). Overall, taking our data and published information together, the S100 family of genes can be considered as reliable prognostic markers of COVID-19 infection and disease progression. Another host factor LCN2, which came up in our study was previously shown to be an important biomarker for viral infection (46, 47), and was also reported to be upregulated in transcriptomic and proteomic studies in COVID-19 patients (48, 49). Furthermore, Serine protease inhibitors (SERPINs) family genes SERPINB3 and SERPINB1 were present among the initially selected 46 upregulated genes. SERPINB3 was at the top of cumulative upregulation ranking (Figure 2C) and in the COVID-19 cohort, it was significantly upregulated in the PS category. It is an inhibitor of papain-like cysteine proteases such as cathepsin, which is required for Spike cleavage during SARS CoV-2 entry (50). Interestingly SERPINA1 deficiencies or mutations in populations were found to be associated with severe forms of COVID-19 (46, 47). Taken together this indicates a potential antiviral role for SERPINs against SARS-CoV-2, which needs further exploration.

Finally, one gene of interest which passed the rigor of meta-analysis was TXN. Although its cumulative upregulation or prognostic values were not very high, we explored its potential as a therapeutic target. Thioredoxin is a small redox protein that plays an active role in keeping the intracellular compartment in a reduced state, which is important to prevent protein aggregation (51, 52). The thioredoxin system consists of three components namely thioredoxin, thioredoxin reductase, and the reducing agent nicotinamide adenine dinucleotide phosphate (NADPH). Thioredoxin reductase is a redox homeostatic enzyme, that can be inhibited by FDA-approved, gold-containing triethyl phosphine drug Auranofin (17, 53). This drug has been shown to have inhibitory activity against rheumatoid arthritis, cancer, HIV/AIDS, parasitic, and bacterial infections (54). A recent study by Rothan *et*.*al* showed Auranofin to inhibit SARS CoV-2 in Huh-7 cells at an EC50 of 1.4µM (18). In comparison, our data in HEK-ACE2 cells showed improved antiviral activity at much lower concentrations of the drug (selectivity index - 33.3, versus 4.07), as evidenced by a decrease in levels of both viral RNA and spike protein expression (Figure 5B and 5C). We went on to validate the antiviral activity of Auranofin in the preclinical hamster challenge model. Results showed a significant reduction in lung viral load and rescue of animal body weight, when administered therapeutically, which may be attributed to the anti-inflammatory activity of the compound (55). Notably, Auranofin has been shown to decrease pro-inflammatory cytokines IL-6, IL1β, and TNFα mRNA levels during SARS-CoV-2 infection in vitro, which are known mediators of disease severity (18). Thioredoxin mRNA levels were upregulated in hamsters, which is consistent with the observation in COVID-19 patients. Auranofin also has inhibitory effects on the PI3K/AKT/mTOR pathway (56), which is required for SARS-CoV-2 viral protein translation (57, 58). This may also contribute to its mechanism of action, however, that needs to be further investigated.

Overall, this study highlights the value of comprehensive analyses of Omics datasets to gain insight into infection biology and identify avenues for potential therapeutic targeting. The selected gene expression data obtained with the COVID-19 cohort reaffirmed the heterogeneity of individual immune response, the role of age, sex, and effect of viral load, all of which are in coherence with observations made by other research groups. We especially uncover the prognostic value of S100 family genes in nasal swabs, many of which are soluble secretory factors and can be easily tested by RT-PCR or ELISA-based methods in nasal swabs to understand the disease progression. Finally, the identification of Auranofin, a safe drug already in clinical use for other medical conditions, as a COVID-19 treatment option culminates the importance of our study and meta-analysis approach in translating virus-host interaction Big Data into clinical interventions.

## STAR⋆METHODS

- KEY RESOURCES TABLE
- RESOURCE AVAILABILITY
  - Lead contact
  - Materials Availability
  - Data and Code Availability
- EXPERIMENTAL MODEL AND SUBJECT DETAILS
  - Ethics Statement
  - Human Subjects
  - Animal Models
  - Cells and Viruses
- METHOD DETAILS
  - OMICs data collection and processing
  - Gene set overlap analysis
  - Gene ontology, Interferome, cellular and tissue localization analysis
  - Virus-Host protein-protein interaction network analysis
  - Nasopharyngeal swab collection and RNA isolation
  - qRT-PCR based measurement of cellular gene expression
  - Cytotoxicity assay
  - Infection in HEK-ACE2 cells
  - Western blot
  - Animal experiment
  - qRT-PCR for viral RNA copy number calculation
- QUANTIFICATION AND STATISTICAL ANALYSIS

## ACKNOWLEDGEMENTS

This study was supported by funds from the DBT-IISc partnership program (DBT (IED/4/2020-MED/DBT)) and the Infosys Young Investigator award (YI/2019/1106) in the ST lab. ST is an Intermediate fellow of Wellcome trust-DBT India Alliance (IA/I/18/1/503613). AS is a Senior fellow of Wellcome trust-DBT India Alliance (IA/S/16/2/50270). AB is supported by KVPY (Kishore Vaigyanik Protsahan Yojana) fellowship from DST, India. SS is supported by PMRF (Prime Minister’s Research Fellowship) from the Ministry of Education, India. We thank Prof. Umesh Varshney and Prof. K.N.Balaji for their administrative guidance.

## AUTHOR CONTRIBUTIONS

ST conceived the study. AB, OK, RN, RR performed the experiments. ST, AB, OK, RN, SS, RS, DS, DG analyzed the data. SM, HB, MJ, DS, AS provided patient samples. AB, SS, ST wrote the manuscript.

## STAR⋆METHODS

## KEY RESOURCES TABLE

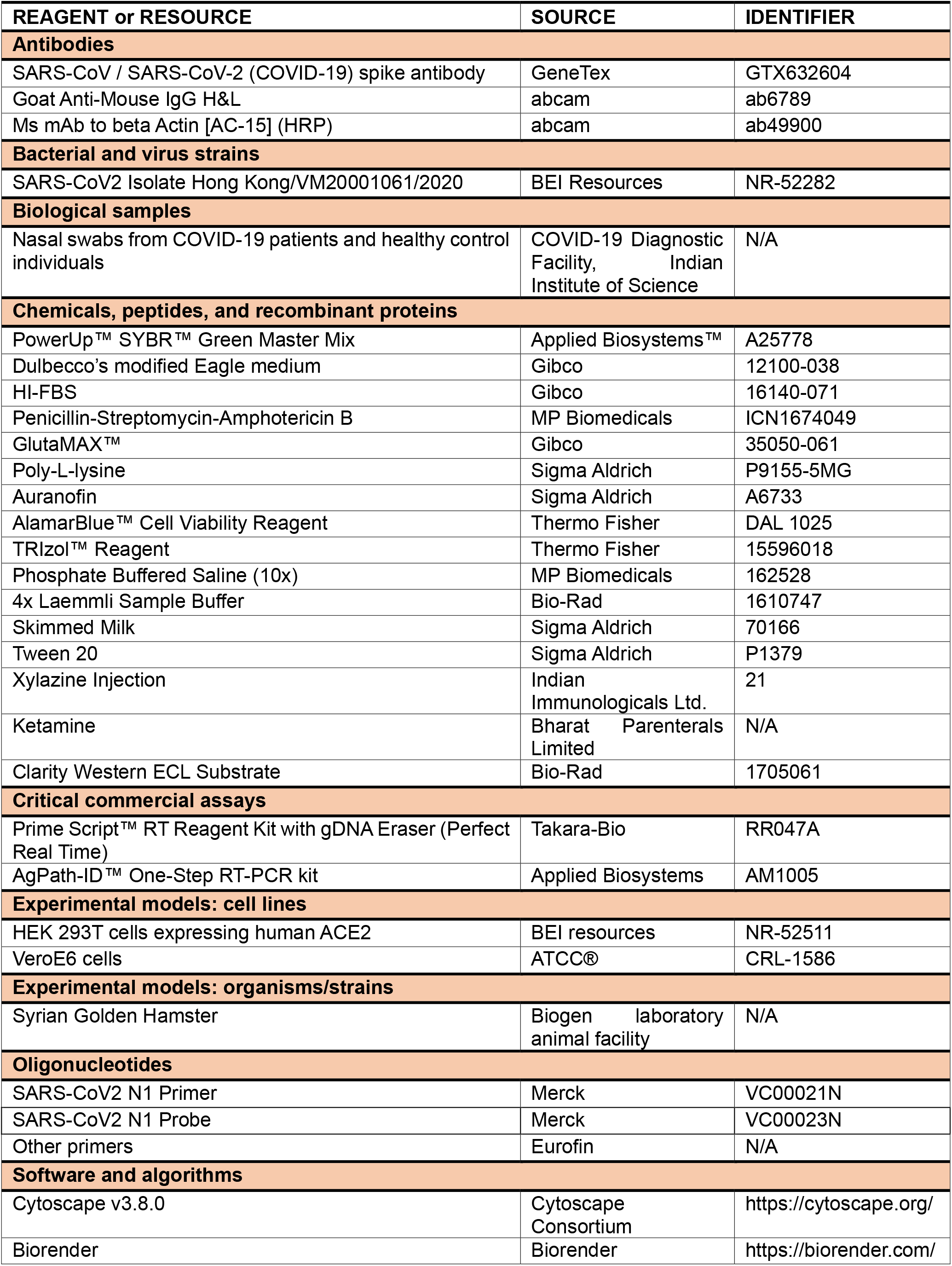
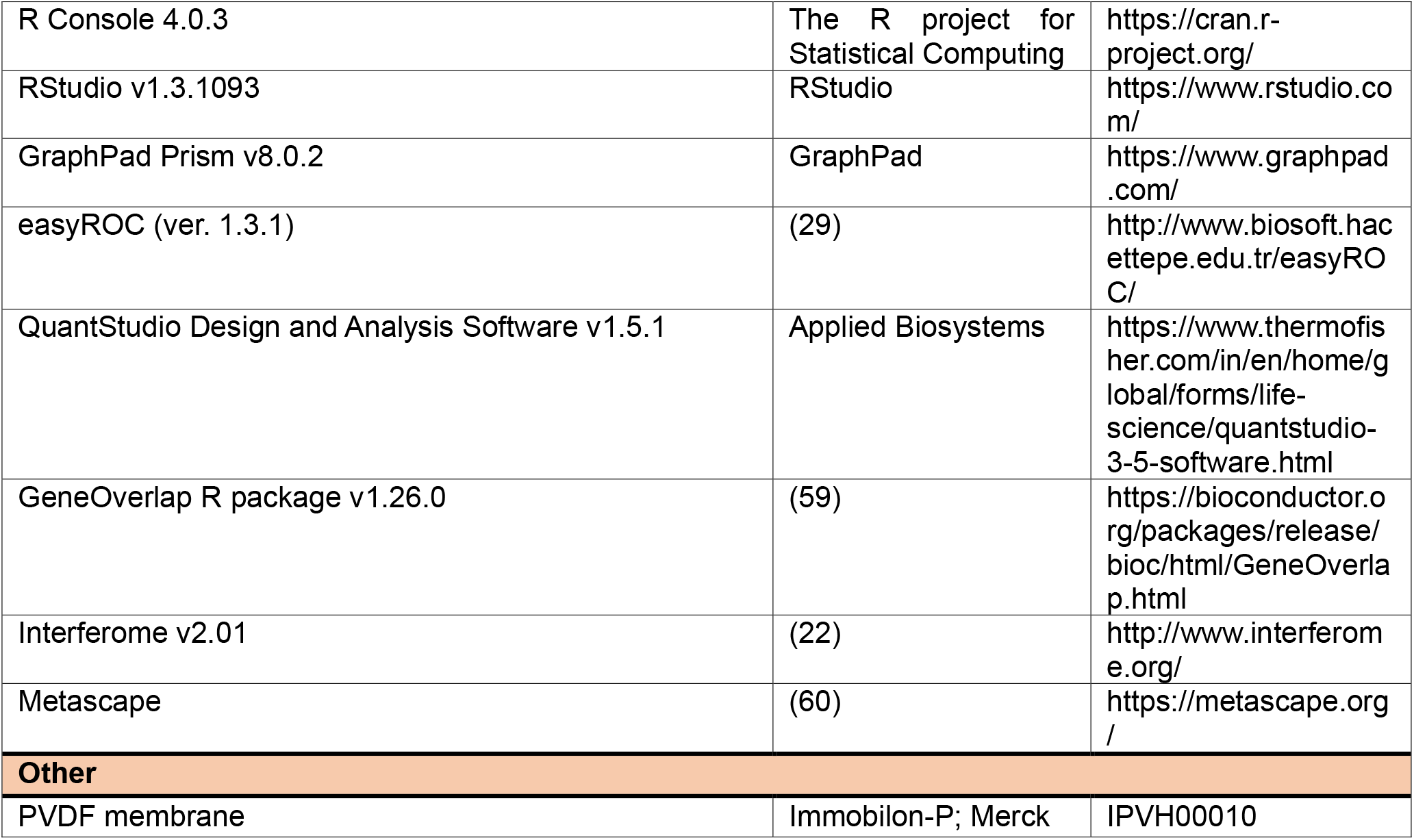

| REAGENT or RESOURCE | SOURCE | IDENTIFIER |
| --- | --- | --- |
| **Antibodies** |  |  |
| SARS-CoV / SARS-CoV-2 (COVID-19) spike antibody | GeneTex | GTX632604 |
| Goat Anti-Mouse IgG H&L | abcam | ab6789 |
| Ms mAb to beta Actin [AC-15] (HRP) | abcam | ab49900 |
| **Bacterial and virus strains** |  |  |
| SARS-CoV2 Isolate Hong Kong/VM20001061/2020 | BEI Resources | NR-52282 |
| **Biological samples** |  |  |
| Nasal swabs from COVID-19 patients and healthy control individuals | COVID-19 Diagnostic Facility, Indian Institute of Science | N/A |
| **Chemicals, peptides, and recombinant proteins** |  |  |
| PowerUp™ SYBR™ Green Master Mix | Applied Biosystems™ | A25778 |
| Dulbecco's modified Eagle medium | Gibco | 12100-038 |
| HI-FBS | Gibco | 16140-071 |
| Penicillin-Streptomycin-Amphotericin B | MP Biomedicals | ICN1674049 |
| GlutaMAX™ | Gibco | 35050-061 |
| Poly-L-lysine | Sigma Aldrich | P9155-5MG |
| Auranofin | Sigma Aldrich | A6733 |
| AlamarBlue™ Cell Viability Reagent | Thermo Fisher | DAL 1025 |
| TRIzol™ Reagent | Thermo Fisher | 15596018 |
| Phosphate Buffered Saline (10x) | MP Biomedicals | 162528 |
| 4x Laemmli Sample Buffer | Bio-Rad | 1610747 |
| Skimmed Milk | Sigma Aldrich | 70166 |
| Tween 20 | Sigma Aldrich | P1379 |
| Xylazine Injection | Indian Immunologicals Ltd. | 21 |
| Ketamine | Bharat Parenterals Limited | N/A |
| Clarity Western ECL Substrate | Bio-Rad | 1705061 |
| **Critical commercial assays** |  |  |
| Prime Script™ RT Reagent Kit with gDNA Eraser (Perfect Real Time) | Takara-Bio | RR047A |
| AgPath-ID™ One-Step RT-PCR kit | Applied Biosystems | AM1005 |
| **Experimental models: cell lines** |  |  |
| HEK 293T cells expressing human ACE2 | BEI resources | NR-52511 |
| VeroE6 cells | ATCC® | CRL-1586 |
| **Experimental models: organisms/strains** |  |  |
| Syrian Golden Hamster | Biogen laboratory animal facility | N/A |
| **Oligonucleotides** |  |  |
| SARS-CoV2 N1 Primer | Merck | VC00021N |
| SARS-CoV2 N1 Probe | Merck | VC00023N |
| Other primers | Eurofin | N/A |
| **Software and algorithms** |  |  |
| Cytoscape v3.8.0 | Cytoscape Consortium | https://cytoscape.org/ |
| Biorender | Biorender | https://biorender.com/ |
| R Console 4.0.3 | The R project for Statistical Computing | https://cran.r-project.org/ |
| RStudio v1.3.1093 | RStudio | https://www.rstudio.com/ |
| GraphPad Prism v8.0.2 | GraphPad | https://www.graphpad.com/ |
| easyROC (ver. 1.3.1) | (29) | http://www.biosoft.hacettepe.edu.tr/easyROC/ |
| QuantStudio Design and Analysis Software v1.5.1 | Applied Biosystems | https://www.thermofisher.com/in/en/home/global/forms/life-science/quantstudio-3-5-software.html |
| GeneOverlap R package v1.26.0 | (59) | https://bioconductor.org/packages/release/bioc/html/GeneOverlap.html |
| Interferome v2.01 | (22) | http://www.interferome.org/ |
| Metascape | (60) | https://metascape.org/ |
| **Other** |  |  |
| PVDF membrane | Immobilon-P; Merck | IPVH00010 |

## RESOURCE AVAILABILITY

### Lead contact

Further information and requests for resources and reagents should be directed to and will be fulfilled by the lead contact, Shashank Tripathi.

### Materials Availability

This study did not generate new unique reagents.

### Data and Code Availability

The published article includes all data generated or analyzed during this study. No new code was developed for this study.

## EXPERIMENTAL MODEL AND SUBJECT DETAILS

### Ethics Statement

This study was conducted in compliance with institutional human ethics and biosafety guidelines, (IHEC No. 13-11092020; IBSC/IISc/ST/17/2020), following the Indian Council of Medical Research and Department of Biotechnology recommendations. All experiments involving animals were reviewed and approved by the Institutional Animal Ethics Committee (Ref: IAEC/IISc/ST/784/2020) at the Indian Institute of Science and conducted in Viral Biosafety level-3 facility.

### Human Subjects

Nasopharyngeal swabs were collected from COVID-19 patients and healthy individuals for diagnostic purposes by hospitals from Bengaluru Urban city and brought to COVID-19 Diagnostic Facility at the Indian Institute of Science in viral transport media (VTM). RNA from patients was isolated using kits recommended and provided by the Indian Council of Medical Research. Samples were chosen to have an almost equal number of patients falling into categories of age, sex, COVID-19 status, and symptomatic status (Table 1). Demographic information was not used as an inclusion criterion.

### Animal Models

All animal experiments were performed using 10 to 12-week-old male and female Syrian golden hamsters purchased from Biogen Laboratory Animal Facility (Karnataka, India). They were given access to pellet feed and water *ad libitum*. Males and females were housed separately and maintained on a 12-hour day/night light cycle at the Viral Biosafety level-3 facility at the Indian Institute of Science. Hamsters were euthanized by an overdose of Ketamine (Bharat Parenterals Limited) and Xylazine (21, Indian Immunologicals Ltd).

### Cells and Viruses

HEK 293T cells expressing human ACE2 (NR-52511, BEI resources) and VeroE6 cells (CRL-1586, ATCC®) were cultured in Dulbecco’s modified Eagle medium (12100-038, Gibco) with 10% HI-FBS (16140-071, Gibco), 100 IU/ml Penicillin, 100 μg/ml Streptomycin and 0.25μg/ml Amphotericin-B (Penicillin-Streptomycin-Amphotericin B, ICN1674049, MP Biomedicals) supplemented with GlutaMAX^™^(35050-061, Gibco). SARS-CoV2 (Isolate Hong Kong/VM20001061/2020, NR-52282, BEI Resources) was propagated and titered by plaque assay in Vero E6 cells as described before (61).

## METHOD DETAILS

### Omics Data collection and Processing

Transcriptomics and protein abundance data from COVID-19 patient’s naso- and oropharyngeal swab, bronchoalveolar lavage fluid (BALF), and other respiratory specimens were chosen from PubMed, BioRxiv, and MedRxiv using different combinations of keywords like “**COVID-19, SARS-CoV-2, Transcriptomics, Proteomics, BALF, swab**”. Studies dealing with gene expression profiles of SARS-CoV-2 infected non-human cell lines and tissues were not considered. The SARS-CoV-2 and COVID-19 collections in the EMBL-EBI PRIDE proteomics database (62) were retrieved and used without any modification. In the NCBI GEO database (63) the following combination of terms was used to collect relevant datasets: **((covid-19 OR SARS-COV-2) AND gse [entry type]) AND “Homo sapiens”[porgn: _txid9606]**. The retrieved datasets were then filtered by their date of publication to collect the studies published between the 1^st^ of January 2020 and the 15^th^ of September 2020. The filtration of datasets was carried out using two parameters, fold-change, and its significance value. Genes and proteins with a fold-change value of ≥ 1.5 and q-value ≤ 0.05 were chosen for the overlap analysis. The raw p-value was used for filtering in cases where the adjusted p-value was not provided, albeit with a more stringent cut-off of ≤ 0.01. The UniProt IDs in filtered protein abundance datasets were converted to their corresponding primary Gene Symbols using UniProt (64).

### Gene set overlap analysis

The GeneOverlap class of R package “GeneOverlap” (59) was used for testing whether two lists of genes are independent, which is represented as a contingency table, and then Fisher’s exact test was used to find the statistical significance. Genes with less than 0.01 overlap p-value were selected for further analysis. The number of background genes for proteome-proteome pairwise study and the transcriptome-proteome pairwise study was 25,000, i.e., the number of protein-coding genes in Hg19. For the transcriptome-transcriptome overlap study, the number of background genes was taken to be the union of expressed genes in both the datasets considered.

### Gene Ontology, Interferome, cellular and tissue localization analysis

Enriched GO terms were obtained by express analysis on Metascape (60) and plotted using ggplot2 (65). The database Interferome v2.01 (22) was queried using gene symbols for identifying interferon regulated genes (IRGs) in normal samples of the respiratory system from both in vitro and in vivo experiments in humans. For cellular localization, each gene was queried on UniProt annotation (66) and Human Protein Atlas ver20.0 (67, 68) and then manually annotated. The single-cell expression data of transcripts was also obtained from Human Protein Atlas ver20.0 (Available from http://www.proteinatlas.org/). They were further filtered to obtain cells that are associated with the immune system or respiratory tract.

### Virus-Host protein-protein interaction network analysis

The interaction data for the selected 46 genes were retrieved from publicly available interaction datasets (13). The retrieved information was then used to generate a network map. Cytoscape v3.8.0 (69) was used to construct the interaction network for virus-host protein-protein interaction. STRING database within the Cytoscape store was used to query the proteins to elucidate the interactions between the proteins significantly altered during SARS-CoV-2 infection. The resulting STRING interaction network (confidence ≥0.999 for all the proteins and confidence ≥0.90 for NAMPT; max number of interactors = 10) was merged with the virus-host PPI on Cytoscape.

### qRT-PCR based measurement of cellular gene expression for patient samples

Equal amounts of RNA were converted into cDNA using Prime Script^™^ RT Reagent Kit with gDNA Eraser (Perfect Real Time) (RR047A, Takara-Bio) and then diluted with 80μl nuclease-free water. The gene expression study was conducted using PowerUp^™^ SYBR^™^ Green Master Mix (A25778, Applied Biosystems^™^) with 18srRNA as the internal control and appropriate primers for the genes (Supplementary Table 3).

### Cytotoxicity assay

HEK-ACE2 cells were seeded in a 96-well cell culture dish pre-coated with 0.1mg/mL poly-L-lysine (P9155-5MG, Sigma-Aldrich) and 24hr later, treated with 0, 1, 2, and 4µM Auranofin (A6733, Sigma-Aldrich) in triplicates. Cells were incubated at 37°C, 5% CO2, and 48hr later, cytotoxicity was measured using AlamarBlue^™^Cell Viability Reagent (DAL 1025, Thermo Fisher) as per manufacturer’s instructions.

### Infection in HEK-ACE2 cells

Cells were seeded in a 24-well cell culture dish pre-coated with 0.1mg/mL poly-L-lysine and 24hr later, used for infection. Cells were first pre-treated for 3hr with 0, 0.12, 0.25, 0.5, and 1µM Auranofin in quadruplicates washed once with complete DMEM and subsequently incubated with 0.1 MOI SARS CoV-2 in 100 µl inoculum for 1 hour at 37 C°. Subsequently complete medium restoring the prior dose of the drug was added to the cells. After 48hr, cells were processed separately for western blot analysis and RNA extraction with TRIzol^™^ Reagent (15596018, Thermo Fisher).

### Western Blot

Cells were washed with 1x PBS (162528, MP Biomedicals) and lysed with 1X Laemmli buffer (1610747, BIO-RAD). Cell lysates were loaded and resolved using a 10% SDS-PAGE gel and the separated proteins were transferred onto a PVDF membrane (IPVH00010, Immobilon-P; Merck). Blocking was performed using 5% Skimmed milk (70166, Sigma-Aldrich) in 1xPBS containing 0.05% Tween 20 (P1379, Sigma-Aldrich) (1xPBST) for two hours at room temperature with slow rocking. Primary antibody incubation was performed overnight (12hr) at 4°C using SARS-CoV / SARS-CoV-2 (COVID-19) spike antibody (GTX632604, GeneTex). Secondary antibody incubation was performed for 2 hours at room temperature with slow rocking using Goat Anti-Mouse IgG H&L (ab6789, Abcam). The blots were developed using Clarity Western ECL Substrate (1705061, BIO-RAD).

### Animal Experiments

Toxicity of 1 and 5 mg/kg bodyweight Auranofin was tested on Syrian golden hamsters by oral administration of the drug in 200 µl PBS. The total bodyweight of hamsters was monitored for up to 7 days (Supplementary Fig 6). Infection experiments were performed by intranasal inoculation of animals with 10^5^ PFU SARS-CoV2 in 100µL PBS. The animals were anesthetized using intraperitoneal injections of Ketamine (150mg/kg) (Bharat Parenterals Limited) and Xylazine (10mg/kg) (21, Indian Immunologicals Ltd) cocktail before infection. Prophylactic treatment involved oral administration of Auranofin (5mg/kg/day) 3-, 2-, and 1-day post-infection (dpi) and followed by virus challenge at day 0. The therapeutic treatment regimen used oral administration of Auranofin (5mg/kg/day) starting at 24-hours post-infection (hpi), followed by 2 and 3 dpi. Total body weight was recorded each day during the entire course of the experiment until the animals were sacrificed at 4 dpi. Virus RNA load in lung tissue specimens was detected by q-RT-PCR.

### RT PCR for viral copy number calculation

For qRT-PCR, total RNA was isolated using TRIzol^™^ Reagent (15596018, Thermo Fisher) as per manufacturer’s instruction and equal amounts of RNA was used to determine viral load using AgPath-ID^™^ One-Step RT-PCR kit (AM1005, Applied Biosystems) using primers and probes targeting the SARS CoV-2 N-1 gene (Forward primer: 5’GACCCCAAAATCAGCGAAAT3’ and Reverse primer: 5’ TCTGGTTACTGCCAGTTGAATCTG3’, Probe: (6-FAM / BHQ-1) ACCCCGCATTACGTTTGGTGGACC). Viral copy number was estimated by generating a standard curve using SARS CoV-2 genomic RNA standard.

## QUANTIFICATION AND STATISTICAL ANALYSIS

Statistical analyses and overlaps were performed in the R statistical environment version 4.0.3 via RStudio version 1.3.1093. Plots were made using the ggplot2 package in R (65) and GraphPad Prism v8.0.2. In boxplots, the hinges of boxes represent the first and third quartiles. The whiskers of the boxplot extend to the value which is 1.5 times the distance between the first and third quartiles. Each data point in the boxplot represents one of the triplicates in qRT-PCR for a particular gene in a particular patient sample. Heatmaps were generated using the R package ComplexHeatmap (70). Receiver Operating Characteristic (ROC) curve analysis and Optimal cut-off determination were performed using the online tool easyROC (ver. 1.3.1) (29).

