## Supplementary Information for "Identification of COVID-19 prognostic markers and therapeutic targets through meta-analysis and validation of Omics data from nasopharyngeal samples"

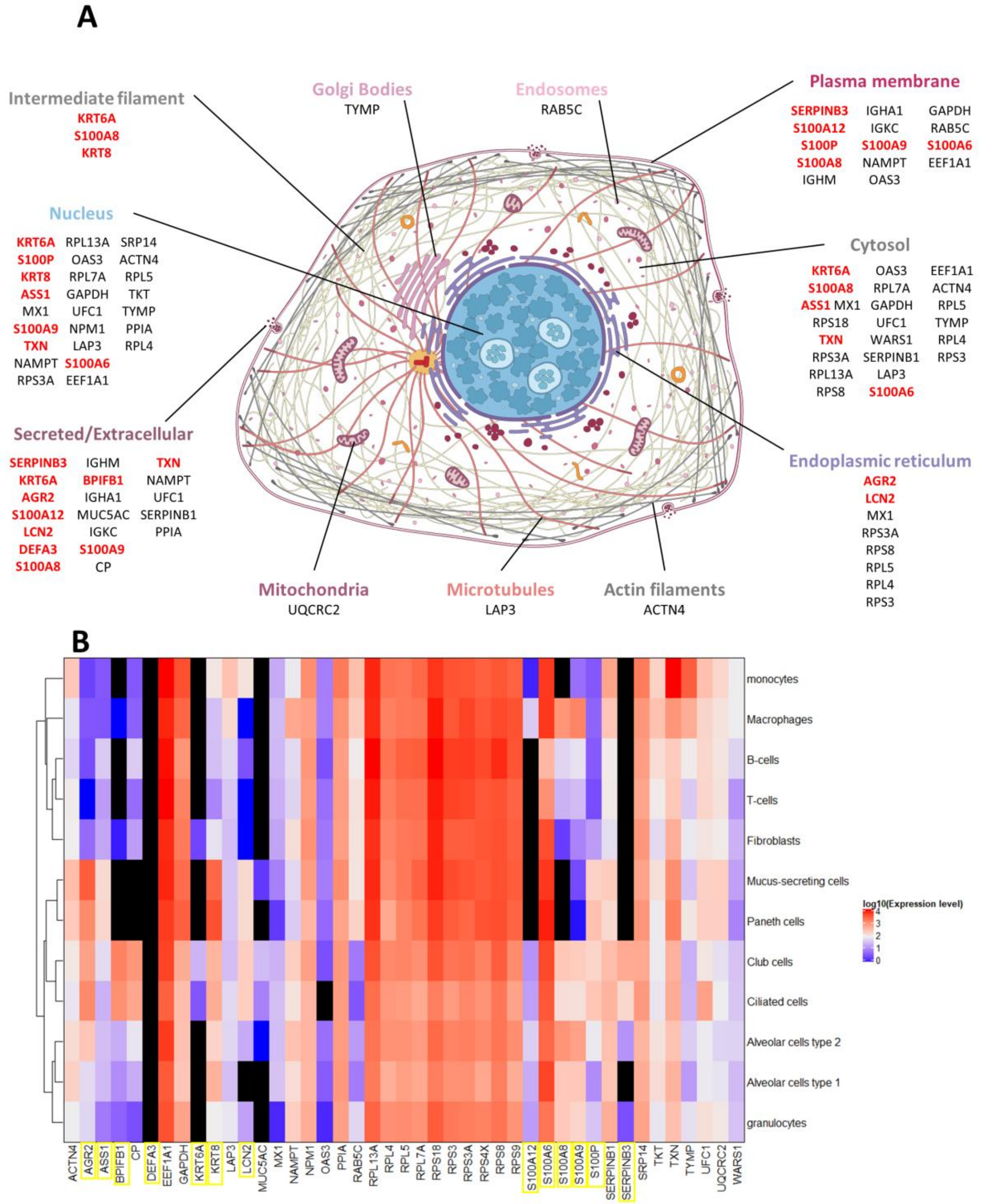


**Supplementary Figure 1: Cellular compartment and cell type-specific localization of selected gene set A)** Cellular localization of proteins that were taken for cumulative score ranking in Figure 2B. Locations were obtained from experimentally verified data in Human Protein Atlas and UniProt (Image credit: Human Protein Atlas). **B)** Single-cell level expression value of the genes which were taken for cumulative score ranking in Figure 2B cells associated with the immune system or respiratory tract. Data was obtained from Human Protein Atlas and the expression level denotes normalized expression value. Highlighted genes in both figures indicate genes whose qRT-PCR validation was performed.

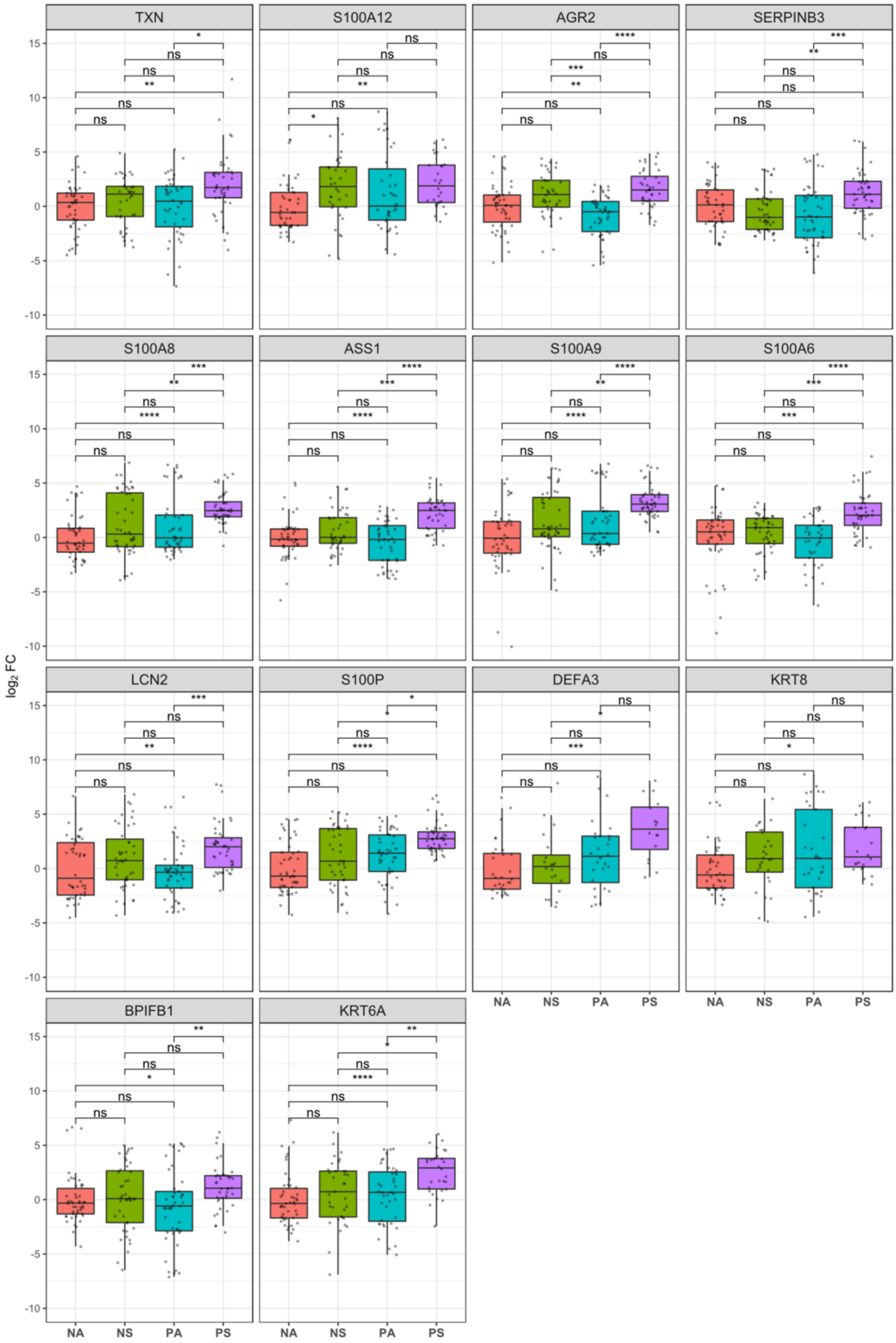
TXN
S100A12
AGR2
SERPINB3
S100A8
ASS1
S100A9
S100A6
LCN2
S100P
DEFA3
KRT8
BPIFB1
KRT6A
$\log_2$ FC
NA
NS
PA
PS

**Supplementary Figure 2: mRNA Expression profile of selected genes in different categories of COVID-19 cohort.** $log_2$ Fold-change values with respect to the NA group are depicted using box plots for each gene without averaging PCR replicates. Differences between Negative asymptomatic (NA), Negative symptomatic (NS), Positive asymptomatic (PA), and Positive symptomatic (PS) groups for each gene were computed using the Kruskal-Wallis test followed by post hoc Dunn's test with Bonferroni corrections for multiple comparisons. *P < 0.05; **P < 0.01; ***P < 0.001; ****P < 0.0001; ns – not significant.

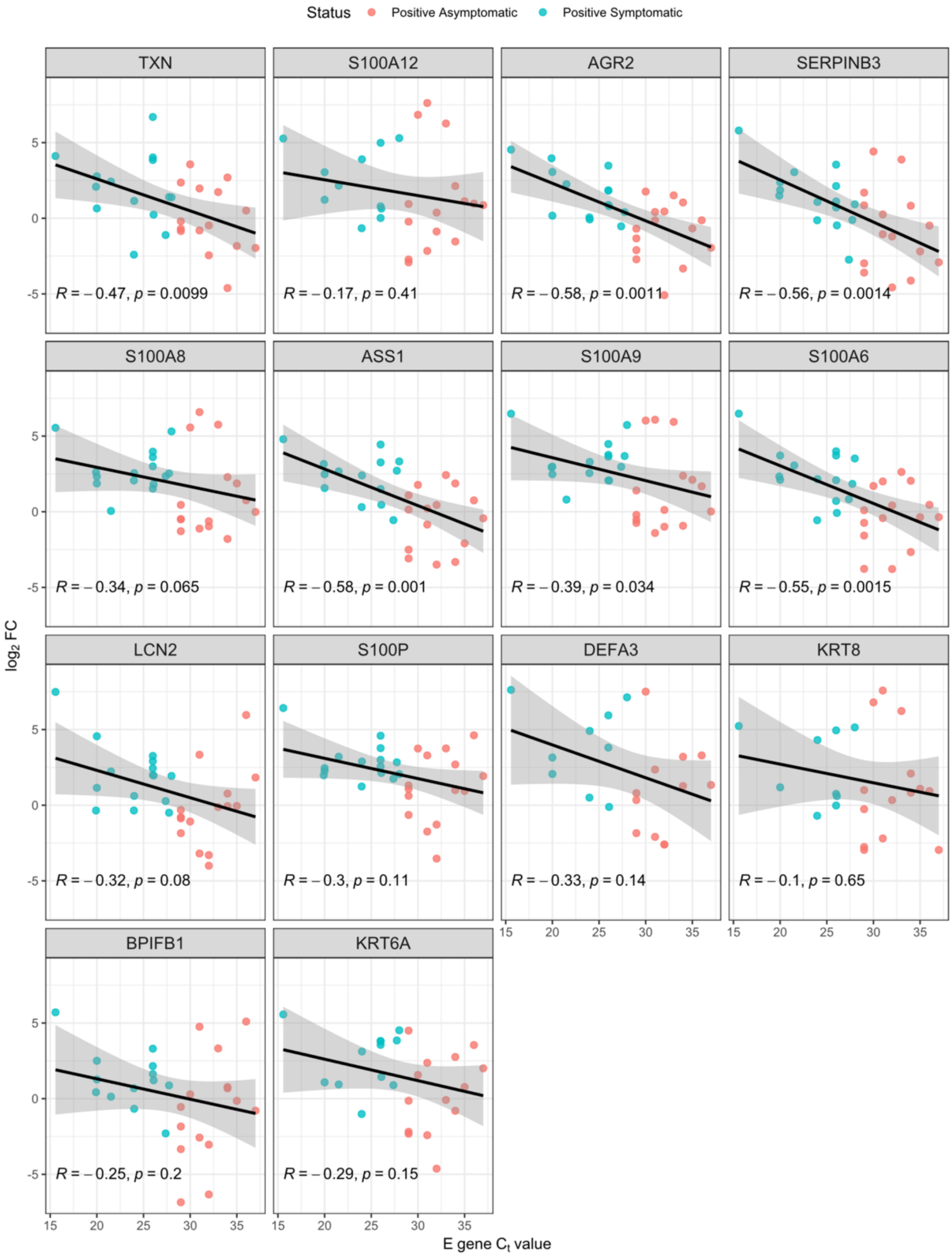

Status
Positive Asymptomatic
Positive Symptomatic
TXN
S100A12
AGR2
SERPINB3
S100A8
ASS1
S100A9
S100A6
LCN2
S100P
DEFA3
KRT8
BPIFB1
KRT6A
R = – 0.47, p = 0.0099
R = – 0.17, p = 0.41
R = – 0.58, p = 0.0011
R = – 0.56, p = 0.0014
R = – 0.34, p = 0.065
R = – 0.58, p = 0.001
R = – 0.39, p = 0.034
R = – 0.55, p = 0.0015
R = – 0.32, p = 0.08
R = – 0.3, p = 0.11
R = – 0.33, p = 0.14
R = – 0.1, p = 0.65
R = – 0.25, p = 0.2
R = – 0.29, p = 0.15
log2 FC
E gene Ct value


39

**Supplementary Figure 3: Correlation of selected gene expression level with viral load in COVID-19 positive patients**. Spearman's correlation (R is the correlation coefficient, and p is the corresponding p-value) $\log_2$ Fold-change values of each gene and $C_t$ value for the viral gene encoding Envelope (E) protein. Each dot represents a patient.

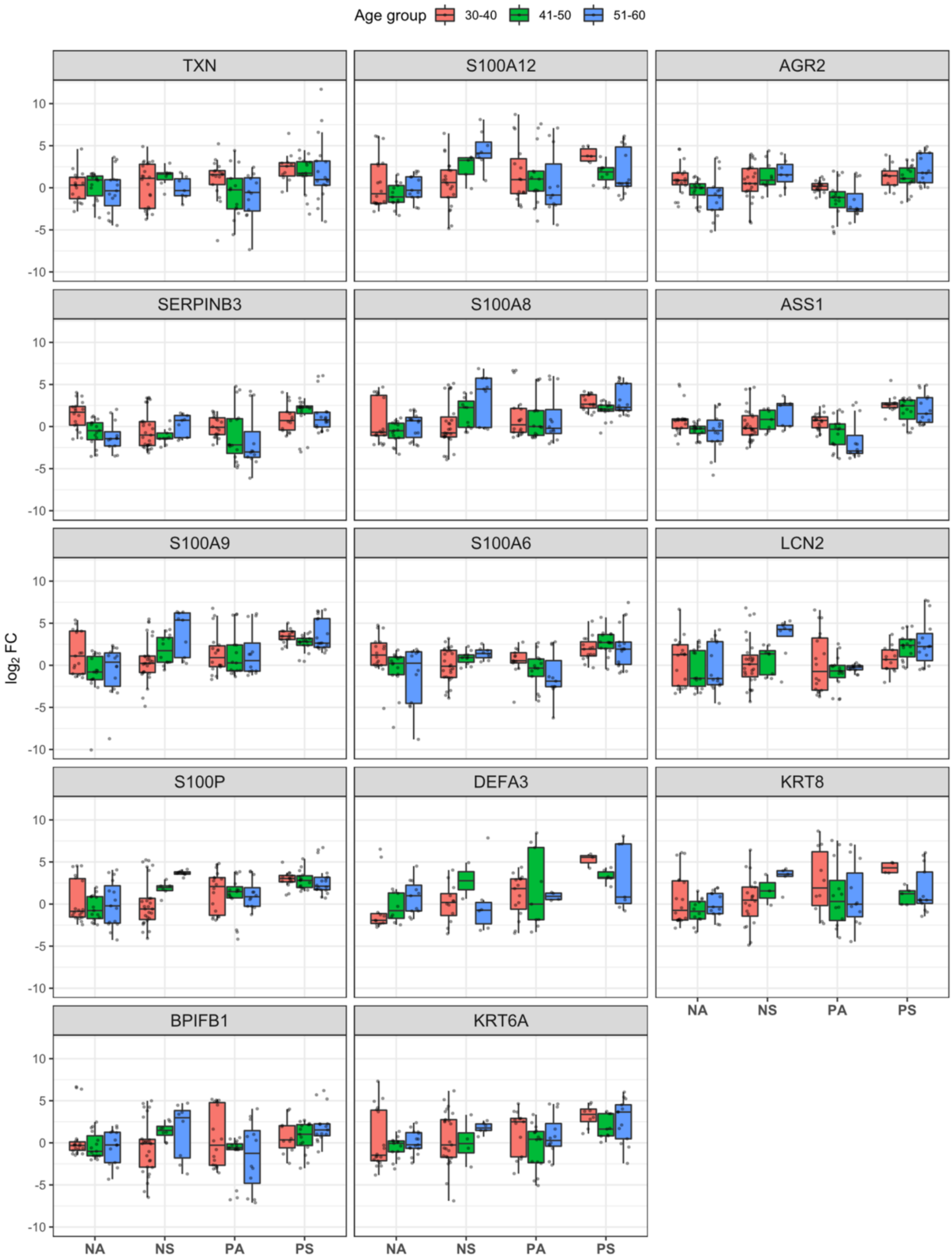


**Supplementary Figure 4: Differences in selected gene expression levels between different age groups.** $\log_2$ Fold-change values with respect to the NA group is depicted using box plots for each gene without averaging PCR replicates for patient groups (Negative asymptomatic (NA), Negative symptomatic (NS), Positive asymptomatic (PA), and Positive symptomatic (PS)) belonging to different age groups.

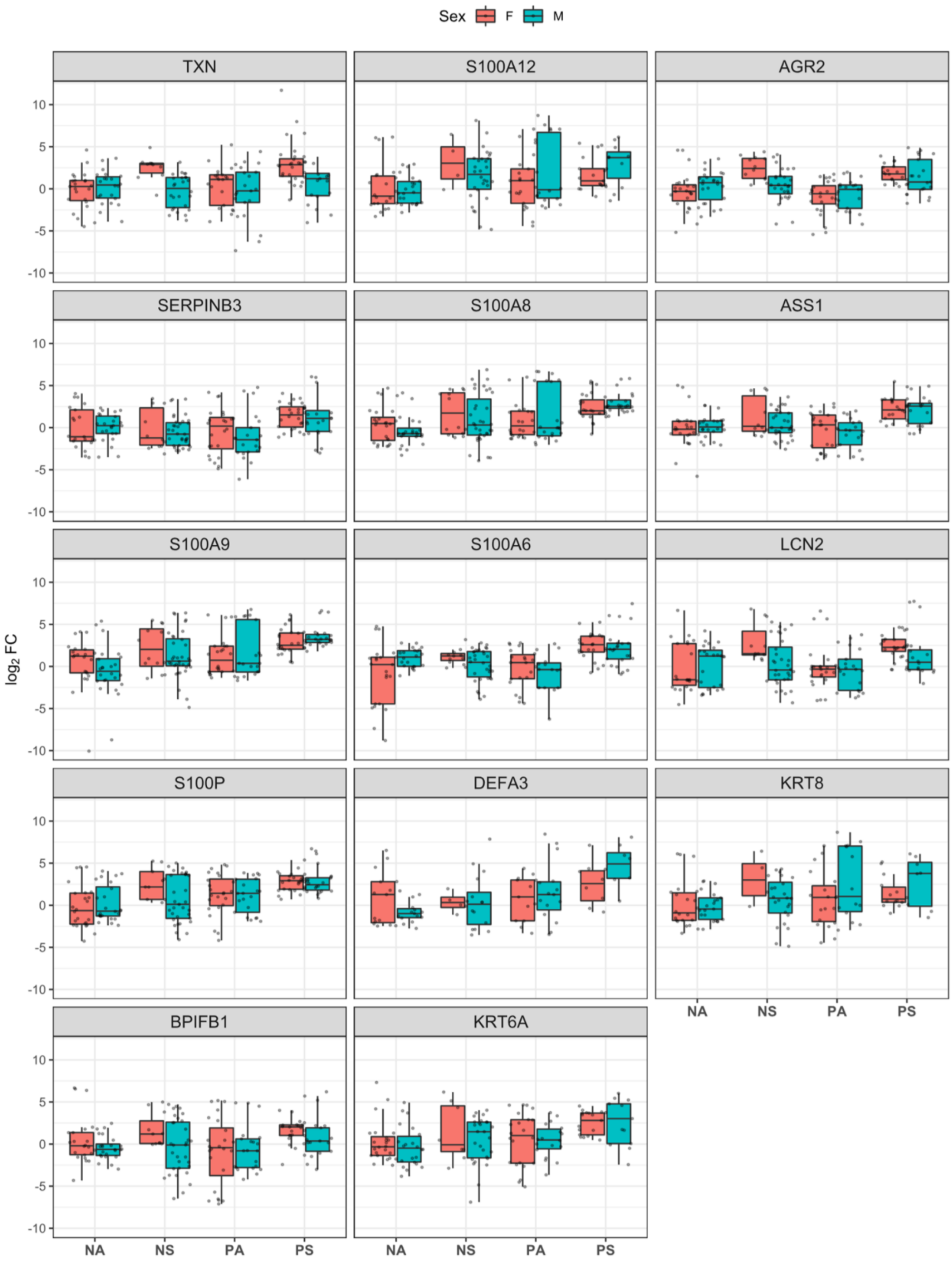


**Supplementary Figure 5: Differences in selected gene expression levels between Male and Female individuals.** $\log_2$ Fold-change values are plotted for patient groups Negative asymptomatic (NA), Negative symptomatic (NS), Positive asymptomatic (PA), and Positive symptomatic (PS) without averaging PCR replicates.

**A**

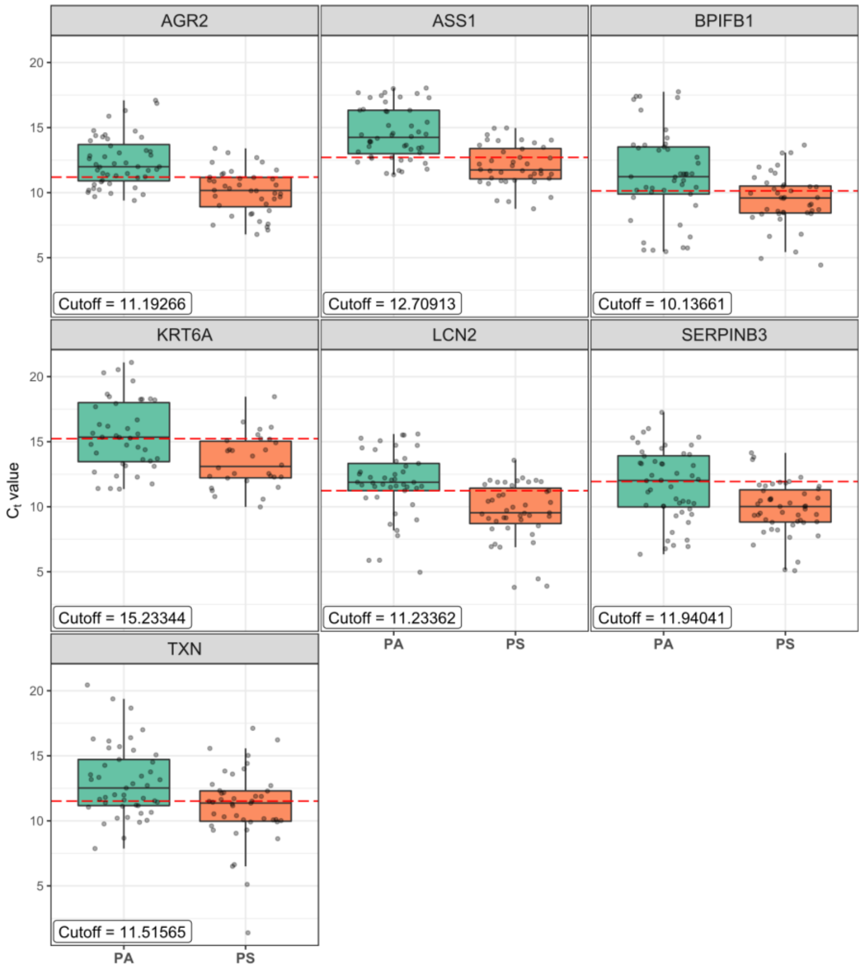


**B**

| Marker | Optimal $C_t$ Cut-off determined by ROC01 | Sensitivity | Specificity |
|---|---|---|---|
| LCN2 | 11.23362 | 0.744 | 0.756 |
| AGR2 | 11.19266 | 0.775 | 0.708 |
| ASS1 | 12.70913 | 0.7 | 0.771 |
| SERPINB3 | 11.94041 | 0.915 | 0.521 |
| BPIFB1 | 10.13661 | 0.65 | 0.711 |
| KRT6A | 15.23344 | 0.833 | 0.550 |
| TXN | 11.51565 | 0.581 | 0.667 |

**Supplementary Figure 6: Optimal $C_t$ cut-offs, Sensitivity, and Specificity of genes after ROC curve analysis. A)** Boxplot of Ct values for Positive asymptomatic (PA) and Positive symptomatic (PS) patients. The red dashed line shows the optimal Ct cut-off determined by the ROC01 method (also shown in the label in each graph). **B)** Optimal Ct cut-off, sensitivity, and specificity values of genes.

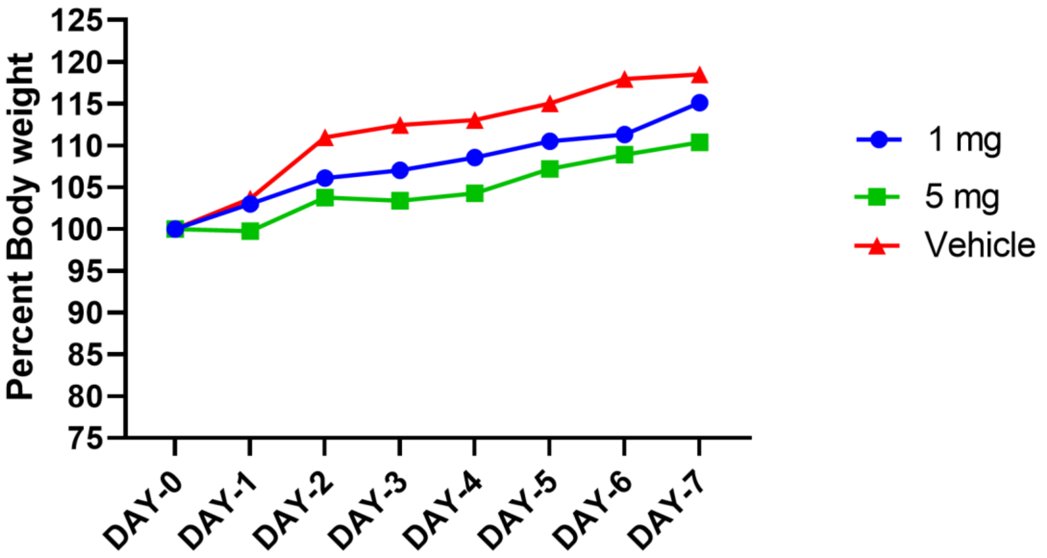


**Supplementary Figure 7: Auranofin toxicity in uninfected hamsters:** 10–12-week-old male and female Hamsters (n=3 per group) were orally administered 200 µl vehicle (DMSO+PBS) or the indicated amount of drug (1 mg or 5 mg) per kg body weight on 3 consecutive days, followed by which body weight was monitored up to Day 7. Percent weight was plotted on the graph considering weight on D0 as 100%.

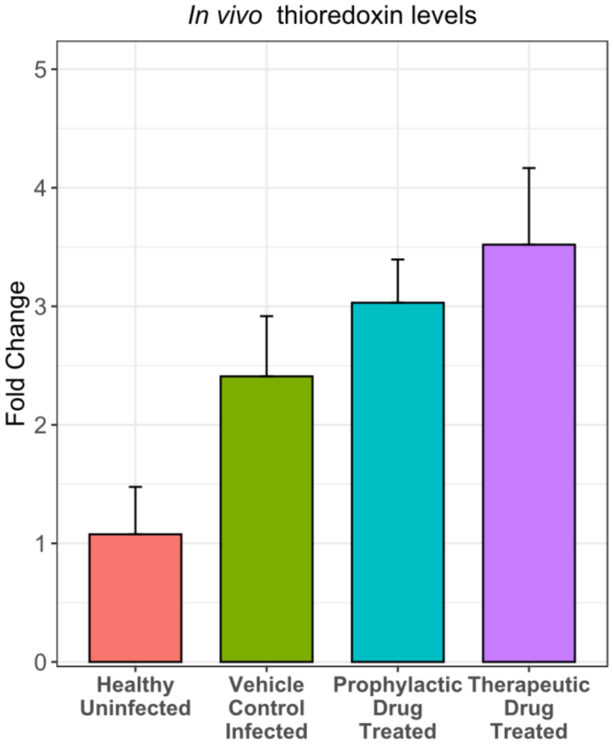


**Supplementary Figure 8: TXN expression in the lungs of SARS-CoV-2 infected hamsters:** From the experiment described in figure 5c, TXN expression in the RNA from lung tissue was measured by qRT-PCR. The fold change in TXN expression as compared to the healthy control group is plotted in the graph. Each column indicates data from 4 animals and error bars indicate mean + standard error.

| Study Type | Study ID | Tissue type | Description | Reference |
|---|---|---|---|---|
| Transcriptomics | T1 | Nasopharyngeal Swabs | Patients 430, Healthy Controls 54 | (1) |
| | T2 | BALF | Patients 86, Healthy Controls 5 | (2) |
| | T3 | BALF | Patients 2, Healthy Controls 3 (from the previous study) | (3) |
| | T4 | BALF | Patients 8, Healthy Controls 20 | (4) |
| Proteomics | P1 | Nasopharyngeal Swabs | 5 positive and five negatives oro- and nasopharyngeal swabs | (5) |
| | P2 | Nasopharyngeal and oropharyngeal swabs | 15 PCR Positive and 15 PCR Negative Patients | (6) |
| | P3 | Nasopharyngeal and oropharyngeal swabs | 20 Positive and 20 Negative Patients | (7) |

**Supplementary Table 1. Overview of studies from where datasets were chosen for Meta-analysis and validation.** The papers discussing proteome and transcriptome of COVID-19 patients were collected from PubMed, MedRxiv, and BioRxiv using combinations of keywords and selected based on the patient sample used and date of publication (materials and methods).

| Study Type | Study ID | No. of genes after filtering | Filter applied |
|---|---|---|---|
| Transcriptomics | T1 | 25 | $log_2FC \geq 0.5849625$, adjusted p value ≤ 0.05 |
| | T2 | 249 | $log_2FC \geq 0.5849625$, adjusted p value ≤ 0.05 |
| | T3 | 2767 | $log_2FC \geq 0.5849625$, adjusted p value ≤ 0.05 |
| | T4 | 1014 | $log_2FC \geq 0.5849625$, adjusted p value ≤ 0.05 |
| Proteomics | P1 | 23 | $log_2FC \geq 0.5849625$, p-value ≤ 0.01 |
| | P2 | 21 | $log_2FC \geq 0.5849625$, q value ≤ 0.05 |
| | P3 | 69 | $log_2FC \geq 0.5849625$, p-value ≤ 0.01 |

**Supplementary Table 2: Number of genes after filtering based on fold-change and statistical significance.** Filtering was performed on DGE data as provided by the authors.

| Sl No | Gene | Organism | Forward Primer (5'-3') | Reverse Primer (5'-3') |
|---|---|---|---|---|
| 1 | TXN | Humans | AGCTCTGTTTGGTGCTTTGG | GCAGTCTTGCTCTCGATCTG |
| 2 | S100A12 | | AGCATCTGGAGGGAATTGTCA | GCAATGGCTACCAGGGATATGAA |
| 3 | S100A8 | | ATGCCGTCTACAGGGATGAC | ACTGAGGACACTCGGTCTCTA |
| 4 | AGR2 | | GTCAGCATTCTTGCTCCTTGT | GGGTCGAGAGTCCTTTGTGTC |
| 5 | SERPINB3 | | CGCGGTCTCGTGCTATCTG | ATCCGAATCCTACTACAGCGG |
| 6 | ASS1 | | CTTGGGGCCAAAAAGGTGTTC | GAGGTAGCGGTCCTCATACAG |
| 7 | S100A6 | | TCCAGAAGGAGCTCACCATT | TCACCTCCTGGTCCTTGTTC |
| 8 | S100A9 | | AAACACTCTGTGTGGCTCCT | TGGTCTCTATGTTGCGTTCCA |
| 9 | LCN2 | | TCACCTCCGTCCTGTTTAGG | CGAAGTCAGCTCCTTGGTTC |
| 10 | KRT6A | | GACCTGGTGGAGGACTTCAA | CTTGGCTTGCAGTTCAACCT |
| 11 | S100P | | AGAAGGAGCTACCAGGCTTC | TTGCAGCCACGAACACTATG |
| 12 | DEFA3 | | CAAAGCATCCAGGCTCAAGG | AGATGCAGGTTCCATAGCGA |
| 13 | KRT8 | | TTAAGGATGCCAACGCCAAG | TTCTGCATCCCAGACTCCAG |
| 14 | 18s rRNA | | GTAACCCGTTGAACCCCATT | CCATCCAATCGGTAGTAGCG |
| 15 | TXN | Hamster | AGCTGATCGAGAGCAAGGAA | CCGAGAAGTCCACGACTACA |

**Supplementary Table 3: Primers used in the study for qRT-PCR.**
